# Regulation of GluA1 Phosphorylation by D-amphetamine and Methylphenidate in the Cerebellum

**DOI:** 10.1101/2020.07.10.196766

**Authors:** Laura Cutando, Emma Puighermanal, Laia Castell, Pauline Tarot, Federica Bertaso, Patricia Bonnavion, Alban de Kerchove d’Exaerde, Elsa Isingrini, Micaela Galante, Glenn Dallerac, Vincent Pascoli, Christian Luscher, Bruno Giros, Emmanuel Valjent

## Abstract

Prescription stimulants, such as d-amphetamine or methylphenidate, are potent dopamine (DA) and norepinephrine (NE) releasers used to treat children and adults diagnosed for attention-deficit/hyperactivity disorder (ADHD). Although increased phosphorylation of the AMPA receptor subunit GluA1 at Ser845 (pS845-GluA1) in the striatum has been identified as an important cellular effector for the actions of these drugs, regulation of this posttranslational modification in the cerebellum has never been recognized. Here, we demonstrate that d-amphetamine and methylphenidate increase pS845-GluA1 in the membrane fraction in both vermis and lateral hemispheres of the mouse cerebellum. This regulation occurs selectively in Bergmann Glia Cells and requires intact norepinephrine release since the effects were abolished in mice lacking the vesicular monoamine transporter-2 selectively in NE neurons. Moreover, d-amphetamine-induced pS845-GluA1 was prevented by β1-adenoreceptor antagonist, whereas the blockade of dopamine D1 receptor had no effect. Additionally, we identified transcriptional alterations of several regulators of the cAMP/PKA pathway, which might account for the absence of pS845-GluA1 desensitization in mice repeatedly exposed to d-amphetamine or methylphenidate. Together, these results point to norepinephrine transmission as a key regulator of GluA1 phosphorylation in Bergmann Glial Cells, which may represent a new target for the treatment of ADHD.

## Introduction

Attention-deficit/hyperactivity disorder (ADHD) is a neurodevelopmental disorder characterized by a myriad of symptoms including distractibility, hyperactivity and impulsivity (Wilens and Spencer, 2010). Children and adults diagnosed for ADHD are generally treated with stimulant medications such as d-amphetamine (Adderrall^®^) and methylphenidate (Ritaline^®^), whose non-medical use and abuse are becoming increasingly prevalent among the general population (Lakhan and Kirchgessner, 2012). Thus, a better understanding of the molecular actions of stimulant exposure on brain function is needed.

The therapeutic efficacy of these drugs as well as their ability to precipitate the development of substance use disorders mainly rely on their actions on dopaminergic and noradrenergic systems (Engert and Pruessner, 2008; Gatley et al., 1996; Pierce and Kalivas, 1997; Sulzer et al., 1995; White and Kalivas, 1998). By rising up extracellular concentrations of dopamine (DA) and norepinephrine (NE) in the striatum and the prefrontal cortex, these drugs increase alertness and attention therefore improving ADHD symptoms (Kuczenski and Segal, 1997) but can also hijack reward processing, motivation, motor and executive functions in case of misuse (Weyandt et al., 2016). Given the widespread DA and NE projections throughout the central nervous system, stimulant-induced molecular adaptations may also exist in other brain regions.

Although the cerebellum has never been considered as a primary site of actions of stimulant medications, compelling evidence indicates that these drugs alter cerebellar neuronal activity as well as monoamine turnover (Freedman and Marwaha, 1980; Quansah et al., 2018; Sorensen et al., 1982) and ameliorate some of the cerebellar abnormalities associated to ADHD symptoms (Stoodley, 2016). The endogenous monoaminergic system is prominent in the cerebellum. Dopaminergic and noradrenergic projections to the cerebellum arise from midbrain DA neurons (VTA/SNc) and hindbrain NE neurons (LC) (Chrapusta et al., 1994; Glaser et al., 2006; Ikai et al., 1994; Ikai et al., 1992; Melchitzky and Lewis, 2000; Nelson et al., 1997; Panagopoulos et al., 1991; Saigal et al., 1980; Saint-Mleux et al., 2004). They modulate the activity of both cerebellar neurons and glial cells, which express a large repertoire of dopaminergic and adrenergic receptors (Juorio et al., 1993; Li et al., 1992; Salm and McCarthy, 1992). However, no work to date has addressed the regulation of intracellular signaling events following stimulants exposure in the cerebellum.

Phosphorylation is a posttranslational modification enabling the rapid functional regulation of proteins. A typical example of such process is the GluA1 subunit of the glutamate α-amino-3-hydroxy-5-methyl-4-isoxazolepropionic acid (AMPA) receptor, which undergoes phosphorylation on two serine residues (pS845 and pS831) achieved by cAMP-dependent protein kinase (PKA) (for pS845) and by calcium/calmodulin-dependent (CaMKII) and protein kinase C (PKC) (for pS831), respectively (Mammen et al., 1997; Roche et al., 1996). Phosphorylation of these two sites directly regulates surface trafficking of GluA1 and its interaction with intracellular proteins as well as electrophysiological properties of AMPA receptors (Roche et al., 1996; Wang et al., 2005). Levels of GluA1 phosphorylation have been reported to be sensitive and modulated by stimulant exposure. For example, acute d-amphetamine and methylphenidate administration increases pS845-GluA1, but not pS831-GluA1, in both the striatum and the prefrontal cortex (Mao et al., 2013; Pascoli et al., 2005; Valjent et al., 2005). AMPA receptors are also widely distributed in the cerebellum (Longone et al., 1998; Martin et al., 1993; Petralia and Wenthold, 1992). However, GluA1 subunit is exclusively expressed by a unique type of astrocyte involved in a wide range of cerebellar functions, the Bergmann Glial cells (BGCs) (De Zeeuw and Hoogland, 2015; Douyard et al., 2007; Saab et al., 2012).

In this study, we probed the ability of d-amphetamine and methylphenidate to modulate GluA1 phosphorylation in BGCs. We also investigated the contribution of DA and NE transmission in the control of stimulants-induced GluA1 phosphorylation and determined the impact of repeated exposure to stimulant medications on these intracellular signaling events.

## Materials and Methods

### Animals

Eight-week-old male C57BL/6 were purchased from Charles River Laboratories. The different transgenic mouse lines used are detailed in Supplementary Table 1. Mice were housed under standardized conditions with a 12 h light/dark cycle, stable temperature (21 ± 2 °C), controlled humidity (55 ± 10%), and food and water ad libitum. All experiments were in accordance with the guidelines of the French Agriculture and Forestry Ministry for handling animals (authorization number/license D34-172-13). Mice were arbitrarily assigned to pharmacological treatments. The number of animals used in each experiment is reported in the figure legends. No statistical methods were used to predetermine sample sizes, but they are comparable to those generally used in the field. For characterization experiments (immunofluorescence, polysome IP, qRT-PCR) male and female mice were used. For Western blot analyses, only male mice were used.

### Drugs

(D)-Methylphenethylamine (D-amphetamine) sulfate salt, methylphenidate hydrochloride, cocaine, GBR12783, propranolol and betaxolol were from Sigma-Aldrich (St. Louis, MO, USA). SCH-23390 was from Tocris (Bristol, UK). All drugs were injected intraperitoneally (i.p.) in a body volume of 10 ml/kg and dissolved in 0.9% (w/v) NaCl (saline) except GBR12783 which was dissolved in H2O. Tamoxifen was dissolved in sunflower oil/ethanol (10:1) to a final concentration of 10 mg/ml and administered i.p. in a volume of 10 ml/kg (100 mg/kg).

### Treatments

Acute pharmacological treatments were carried out with saline, d-amph (10 mg/kg), mph (15 mg/kg), cocaine (20 mg/kg) and GBR12783 (15 mg/kg). Mice were sacrificed 15 min after injections. The various antagonists were injected 30 min prior saline, d-amph (10 mg/kg) or mph (15 mg/kg) injection. Propranolol and betaxolol were administered at 20 mg/kg. SCH-23390 at 0.1mg/kg. For chronic treatments, saline, d-amph (10 mg/kg) and mph (15 mg/kg) were administered once per day during 5 days. Betaxolol (20 mg/kg) was administered 30 min prior saline, d-amph (10 mg/kg) or mph (15 mg/kg) injection. Mice were killed 15 min after the fifth d-amph or mph administration. To induce the Cre expression in the *Gfap-Cre^ERT2^::RiboTag* mice (here referred as *Gfap-RiboTag*), tamoxifen (100 mg/kg) was administered during 3 consecutive days intraperitoneally in a volume of 10 ml/kg.

### Stereotaxic injection in the VTA and the LC

Stereotaxic injections in the VTA of *Slc6a3-Cre* mice have been performed as previously described (Pascoli et al., 2015). Six-week-old C57Bl6 *Dbh-Cre* mice were anaesthetized in isoflurane (4.0% for induction, 0.5–1.0% for maintenance) in oxygen (0.5L/min) and mounted in a stereotaxic frame (Model 940, David Kopf Instruments). The skin on the head was shaved and aseptically prepared, and 2 mg/kg lidocaine infused subcutaneously at the incision site. A single longitudinal midline incision was made from the level of the lateral canthus of the eyes to the lambda skull suture. Injections were performed using a 30-gauge needle (Cooper’s needle Works Ltd) connected by PBS 0.01M-filled tubing to a 10-μL Hamilton syringe in an infusion pump (KDS 310 Plus Nano Legacy Syringe Pump, KD Scientific). Injections were performed at 0.1 μL/min and the needle left in situ for 15 min afterwards to allow diffusion. Injections were performed in LC (AP –5.4 mm from bregma skull suture, ML +/−1.2 mm, DV −3 mm from brain surface, 800 nL). Coordinates based on the mouse brain atlas (Franklin and Paxinos, 2008). Animals received Cre-inducible recombinant AAV vector prepared by the viral vector core at the University of North Carolina (lot AV4311c, 3.6 x 10^12^ genome copies (gc)/mL) AAV2.5-EF1a-DIO-Cherry for anatomical tracing of LC NE neurons projections. Animals were allowed to recover in individual housing after surgery and we awaited at least 8 to 12 weeks for transgene expression in terminals before being killed for histology.

### Tissue preparation and immunofluorescence

Tissue preparation and immunofluorescence were performed as previously described (Biever et al., 2015). Mice were rapidly anaesthetized with Euthasol^®^ (360 mg/kg, i.p., TVM lab, France) and transcardially perfused with 4% (weight/vol) paraformaldehyde in 0.1 M sodium phosphate buffer (pH 7.5). Brains were post-fixed overnight in the same solution and stored at 4°C. Forty-μm thick sections were cut with a vibratome (Leica, France) and stored at −20°C in a solution containing 30% (vol/vol) ethylene glycol, 30% (vol/vol) glycerol and 0.1 M sodium phosphate buffer, until they were processed for immunofluorescence. Sagittal cerebellar sections were identified using a mouse brain atlas (Franklin and Paxinos, 2008) and were processed as follows: free-floating sections were rinsed three times 10 min in Tris-buffered saline (TBS, 50 mM Tris-HCL, 150 mM NaCl, pH 7.5). After 20 min incubation in 0.1% (vol/vol) Triton X-100 in TBS, sections were rinsed in TBS again during 10 min and blocked for 1 h in a solution of 3% BSA in TBS. Cerebellar sections were then incubated 72 hours at 4°C with the primary antibodies (Supplementary Table 2) diluted in a TBS solution containing 1% BSA and 0.15% Triton X-100. Sections were rinsed three times for 10 min in TBS and incubated for 60 min with goat Cy3-coupled anti-rabbit (1:500, Thermo Fisher Scientific Cat# 10520), goat Alexa Fluor 594-coupled anti-chicken (1:400, Thermo Fisher Scientific Cat#A-11042), goat Alexa Fluor 488-coupled anti-chicken (1:500, Thermo Fisher Scientific Cat#A-11039), goat Alexa Fluor 488-coupled anti-mouse (1:500, Thermo Fisher Scientific Cat#A-11001), goat Alexa Fluor 488-coupled anti-rabbit (1:500, Life Technologies Cat#A-11034), goat Cy5-coupled anti-mouse (1:500, Thermo Fisher Scientific Cat#A-10524), goat Cy3-coupled anti-mouse (1:500, Jackson Immunoresearch Cat#115-165-003) or goat Cy5-coupled anti-rabbit (1:500, Thermo Fisher Scientific, Cat#A10523) antibodies. Sections were rinsed for 10 minutes (twice) in TBS and twice in Tris-buffer (0.25 M, pH 7.5) before mounting in DPX (Sigma-Aldrich). Confocal microscopy and image analysis were carried out at the Montpellier RIO Imaging Facility. Images covering the entire cerebellum and double-labeled images from each region of interest were acquired using sequential laser scanning confocal microscopy (Zeiss LSM780). Photomicrographs were obtained with the following band-pass and long-pass filter setting: Alexa fluor 488/Cy2 (band pass filter: 505–530), Cy3 (band pass filter: 560–615) and Cy5 (longpass filter 650). All parameters were held constant for all sections from the same experiment. Three to four slices per mouse were used in all immunofluorescence analyses (n = 2-3 mice / staining).

### Tissue Collection and samples preparation for Western blot analysis

Mice were killed by cervical dislocation and the heads were immersed in liquid nitrogen for 4 sec. The brains were then removed and sectioned on an aluminum block on ice and the whole cerebellum was rapidly isolated from the brainstem. For figures 2a and 2b the whole cerebellum was sonicated in 300 μl of 10% sodium dodecyl sulfate (SDS) and boiled at 95°C for 10 min. For the other figures, subcellular fractionation was performed after tissue collection as previously described (Ozaita et al., 2007). In each experiment, samples from different animal groups, treatments or brain regions were processed in parallel to minimize inter assay variations.

**Figure 1.**
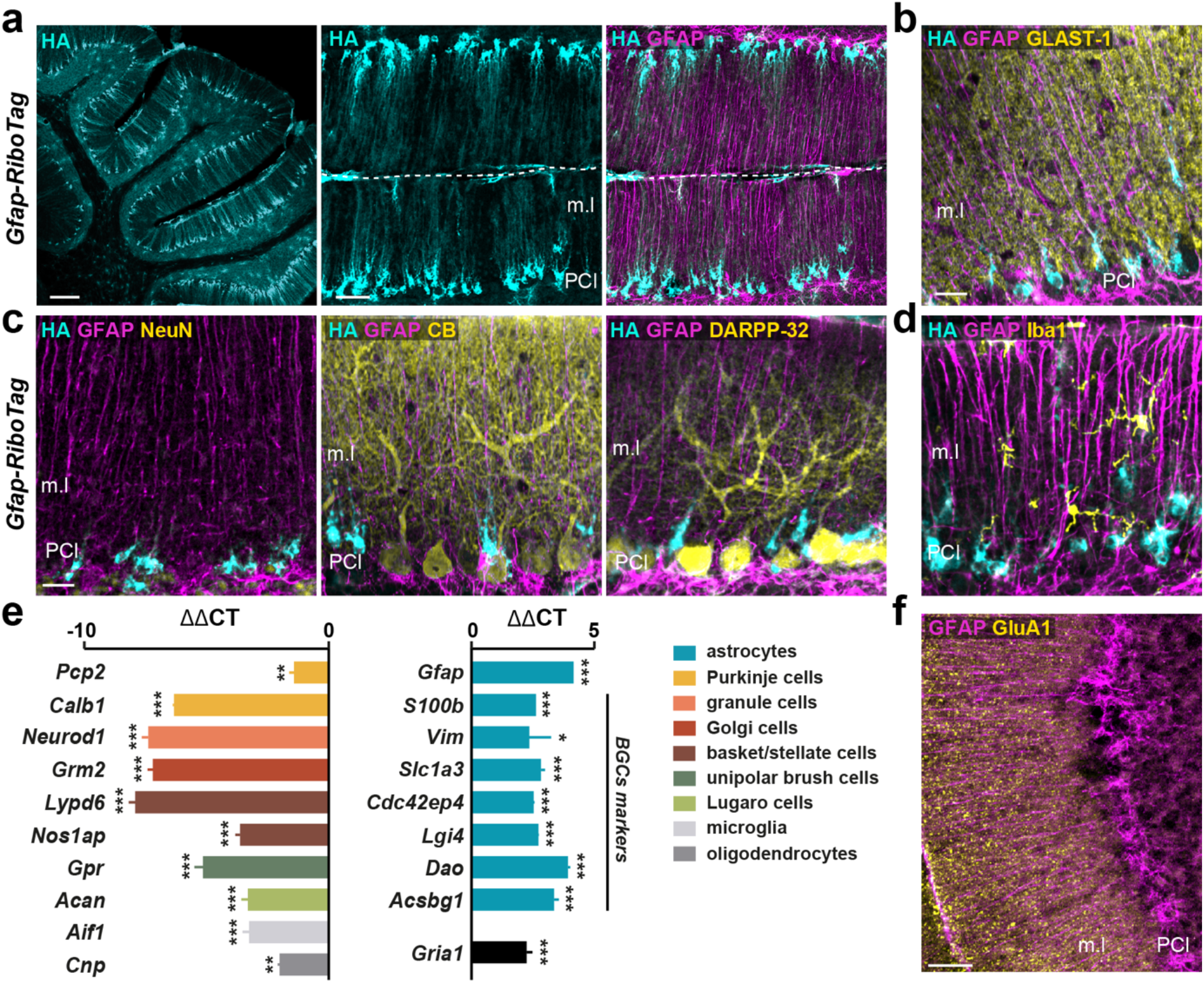
GluA1 are expressed in cerebellar astrocytes in adult mice. (**a**) Sagittal section from *Gfap-RiboTag* mice stained with HA (cyan) showing the distribution of GFAP-expressing cells in the cerebellar cortex. Scale bar: 400 μm (left image). Double immunofluorescence for HA (cyan) and GFAP (magenta). Scale bar: 62 μm (right images). (**b-d**) Triple immunofluorescence for HA (cyan), GFAP (magenta) and the astrocytic marker GLAST-1 (yellow) (**b**), the neuronal markers NeuN, Calbindin-D28k (CB), and DARPP-32 (yellow) (**c**), and the microglial marker Iba1 (**d**). Scale bar: 15 μm. (**e**) Validation by qRT-PCR (ΔΔCT) of the enrichment of glial markers (Cyan) and depletion of transcripts encoding PC, interneurons, oligodendrocyte and microglial markers after HA-immunoprecipitation from cerebellar extract compared with the input fraction (n = 5-6 pooled samples of 2 mice / pool). All genes were normalized to *Tbp2a*. Data were analyzed by two-sided *t* test. \**p* < 0.05; \*\**p* < 0.01; \*\*\**p* < 0.001 (**f**) Double immunofluorescence in the cerebellar cortex for GluA1 (yellow) and GFAP (magenta). Scale bar: 60 μm. Note the expression of GluA1 in BGCs of the m.l. Scale bar: 100 μm. PCl: Purkinje cell layer; g.l: granular layer; m.l: molecular layer. For detailed statistics see Supplementary Table 4.

**Figure 2.**
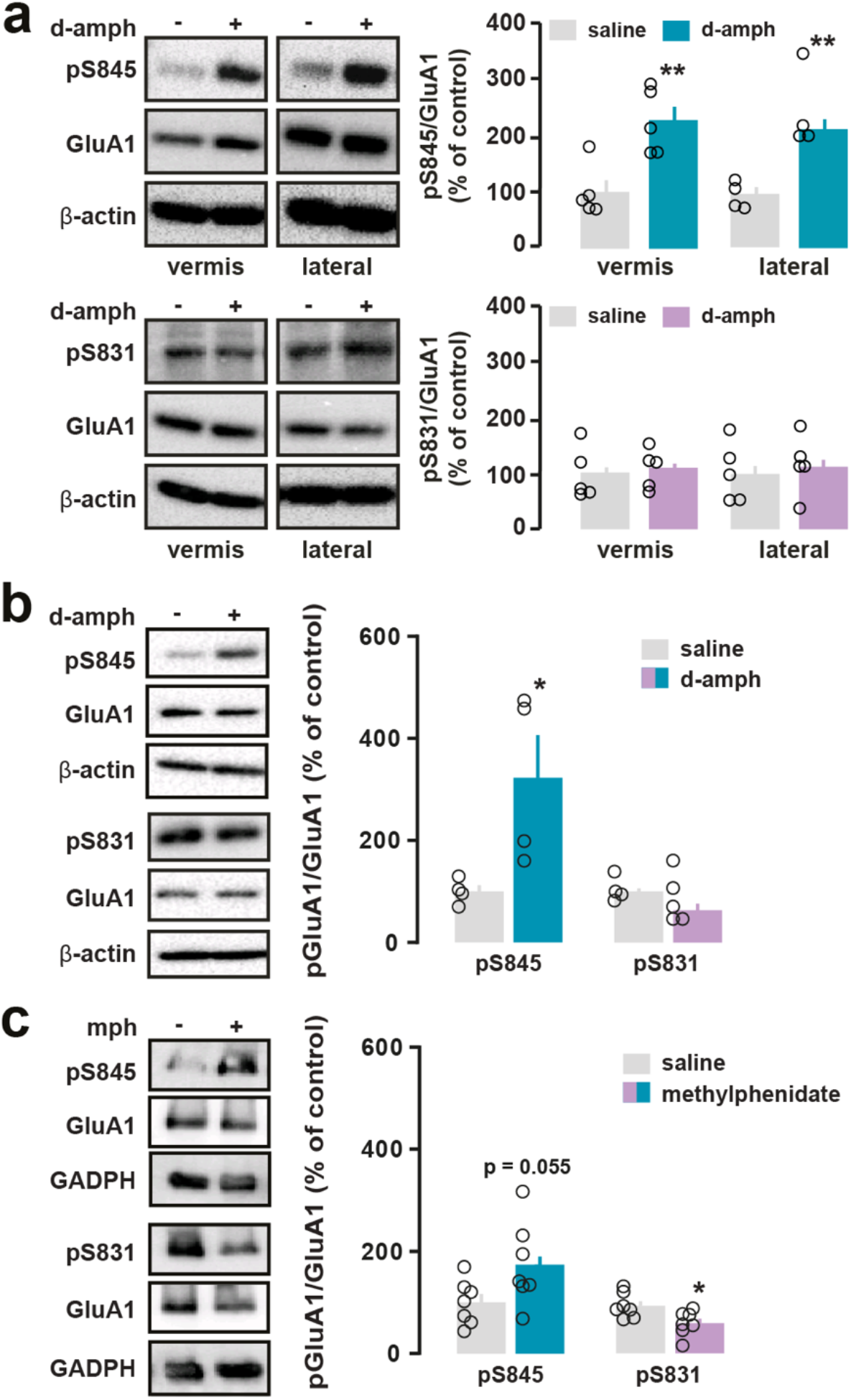
Acute psychostimulant administration increases S845 GluA1 phosphorylation in the cerebellum. (**a**) Representative immunoblots (left) and quantification (right) of GluA1 phosphorylation at S845 (pS845-GluA1) (top) or at S831 (pS831-GluA1) (bottom) in the vermis and lateral hemispheres of C57/Bl6 mice 15 minutes after a single administration of saline or d-amphetamine (d-amph, 10 mg/kg). Phosphorylated forms of GluA1 were normalized to unphosphorylated GluA1. β-actin was used as housekeeping protein. Data are expressed as percentage of saline group (n = 4-5 mice / group for vermis and lateral). Data are presented as means ± SEM and analyzed by two-sided *t* test. \*\**p* < 0.01. (**b-c**) Representative immunoblots (left) and quantification (right) of GluA1 phosphorylation at S845 (pS845-GluA1) (top) or at S831 (pS831-GluA1) (bottom) in membrane cerebellar extracts of C57/Bl6 mice 15 minutes after a single administration of saline, d-amphetamine (d-amph, 10 mg/kg) **(b)** or methylphenidate (mph, 15 mg/kg) (**c**). Phosphorylated forms of GluA1 were normalized to unphosphorylated GluA1. β-actin (**b**) and GAPDH (**c**) were used as housekeeping proteins. Data are expressed as percentage of saline group (n = 4-5 mice / group for d-amph, n = 7 mice / group for mph). Data are presented as means ± SEM and analyzed by two-sided *t* test. \**p* < 0.05. For detailed statistics see Supplementary Table 4.

### Western blot

Protein quantification and western blots were performed following the protocol previously described (Biever et al., 2015). Following the manufacturer’s instructions, protein contents for each sample were determined by BCA protein assay (Pierce) (Lot# RG235624; Thermo Scientific). Equal amounts of cerebellar lysates were mixed with denaturing 4X Laemmli loading buffer. Samples with equal amounts of total protein were separated in 11% SDS-polyacrylamide gel before electrophoretic transfer onto Immobilon-P membranes (#IPVH00010; Millipore). Membranes were cut horizontally, at different molecular weights to be analyzed with different primary antibodies. Using 4% bovine serum albumin (BSA) in 0.1M PBS, membranes were blocked for 45 min, and then incubated for 2h with the primary antibodies (Supplementary Table 2). To detect the primary antibodies, horseradish peroxidase-conjugated antibodies (1:10000) from Cell Signaling Technology to rabbit (Cat# 7074S) or mouse (Cat#7076S) were used, and visualized by enhanced chemiluminescence detection (Luminata Forte Western HRP Substrate; Millipore, Cat# WBWF0500). The optical density of the relevant immunoreactive bands was measured after acquisition on a ChemiDoc Touch Imaging System (Bio-Rad) controlled by Image Lab software version 3.0 (Bio-Rad). Representative cropped immunoblots for display were processed with Adobe Illustrator CS6. For quantitative purposes, the optical density values (O.D) of active phospho-specific antibodies were normalized to the detection of non-phospho-specific antibodies in the same sample and expressed as a percentage of control treatment or group. More specifically, in the Figures 2a, 2c, 4c, 4d, 5g, 6a, since the home-made stripping solution was not strong enough to ensure the removal of the first antibodies tested, the evaluation of pS845-GluA1 or p831-GluA1 and the total form GluA1 was performed in two different gels. The O.D of phospho-sites and the O.D of GluA1 were normalized gel to the O.D of control proteins β-actin or GADPH in each gel. Therefore, the analysis of the O.D was performed as follow: pS845-GluA1 or p831-GluA1/β-actin or GADPH (gel 1) versus GluA1/β-actin or GADPH (gel 2). In Figures 2b and 5d, GluA1 was evaluated after stripping of pS845-GluA1 or p831-GluA1. β-actin was assessed in the same gel. The analysis of the O.D was performed as follow: pS845-GluA1 or p831-GluA1 versus GluA1/β-actin. Finally, in the Figure 6c O.D of pS845-GluA1 was directly normalized to O.D of GluA1. The number of samples and the statistical test used in each experiment are specified in the figure legends.

**Figure 3.**
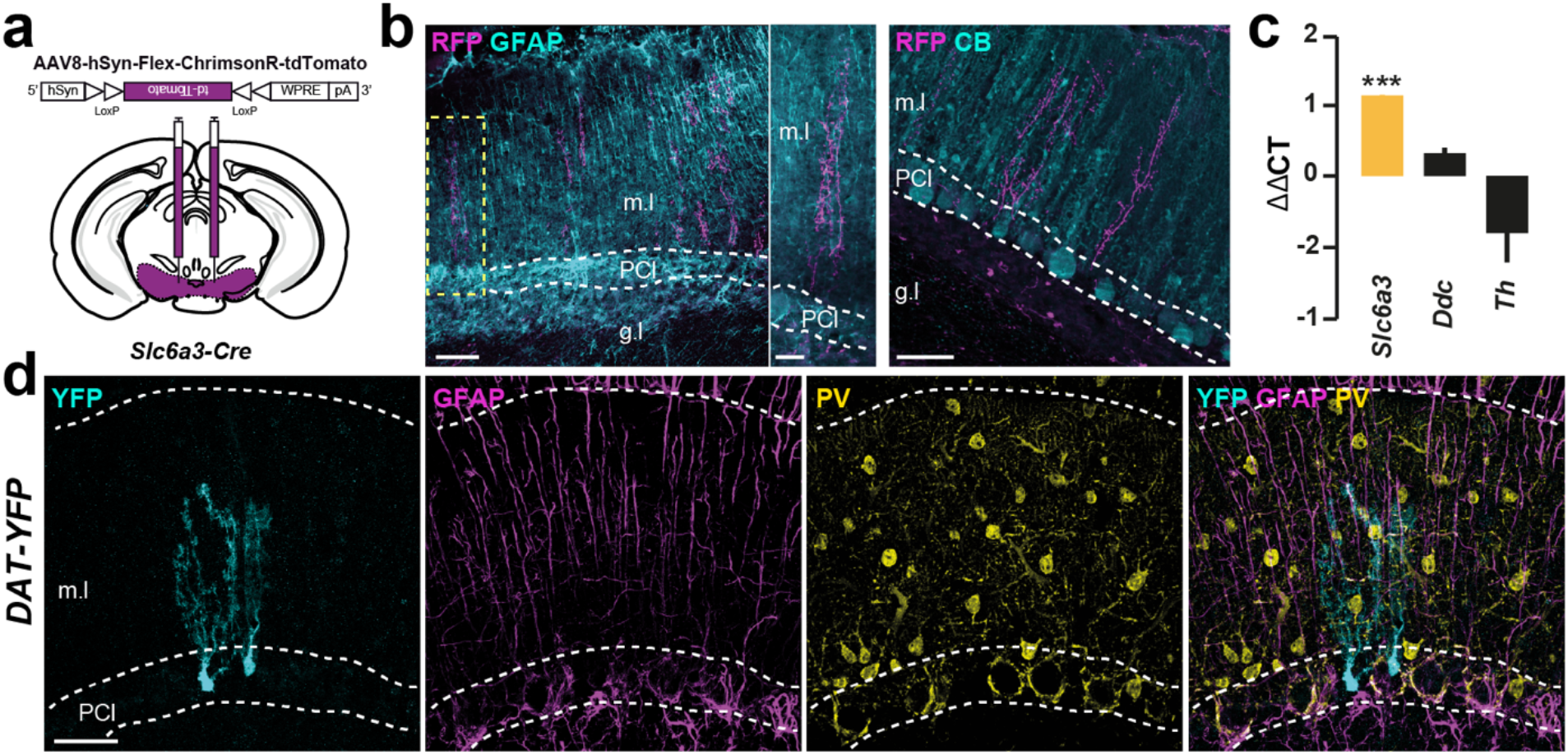
Sources of dopamine in the cerebellar cortex. (**a**) Scheme of Cre-dependent AAV9-CAG-Flex-TdTomato injection in the VTA of *Slc6a3-Cre* mice. (**b**) Double immunofluorescence for RFP (magenta), GFAP (cyan) (left) and calbindin-D28k (CB) (right) identifying RFP-expressing dopamine VTA neuron axons positive fibers in the cerebellar cortex. Note the orientation of RFP-positive fibers paralleling the BGCs. Scale bars: 60 μm (left), 15 μm (middle), 30 μm (right). (**c**) qRT-PCR analysis of *Slc6a3, Th* and *Ddc* transcripts in cerebellar extracts from *Gfap-RiboTag* mice after HA-immunoprecipitation. All genes were normalized to *Tbp2a*. Data are presented as the fold change comparing the pellet fraction versus the input (n = 4-5 pooled samples of 2 mice / pool). Data were analyzed by two-sided *t* test. \*\*\**p* < 0.001. (**d**) Triple immunofluorescence for GFP (cyan), GFAP (magenta) and parvalbumin (PV; yellow) in the cerebellar cortex of *Slc6a3-Cre* mice crossed with the reporter mouse line *ROSA26-YFP* (DAT-YFP). Scale bar: 30 μm. PCl: Purkinje cell layer; g.l: granular layer; m.l: molecular layer. For detailed statistics see Supplementary Table 4.

**Figure 4.**
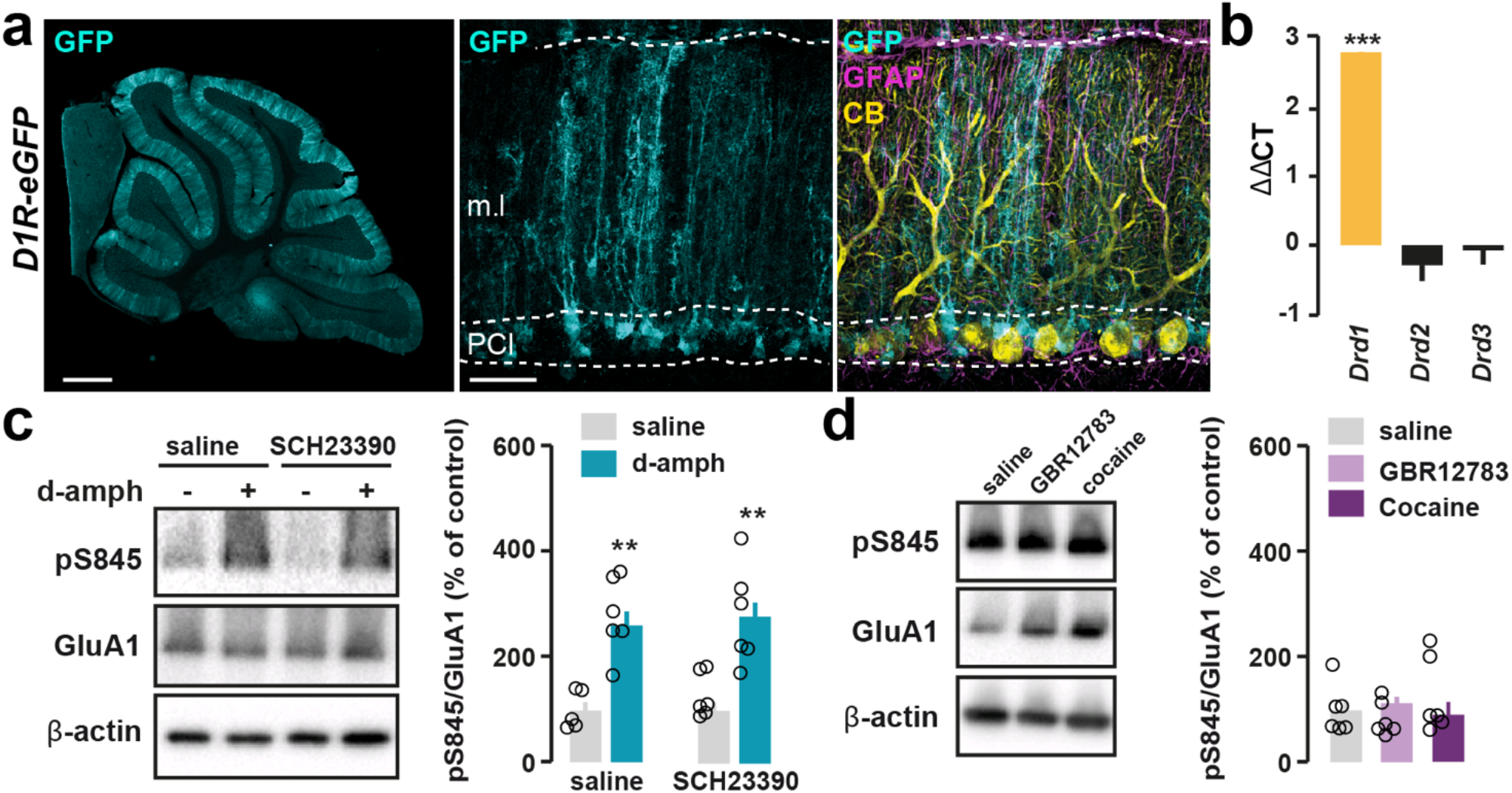
D-amphetamine induced GluA1 phosphorylation in BGCs does not require D1R activation. (**a**) Sagittal section from *D1-eGFP* mice stained with GFP (cyan) showing the distribution of D1R-expressing cells in the cerebellar cortex. Scale bar: 1 mm (left image). Triple immunofluorescence for GFP (cyan), GFAP (magenta), and calbindin-D28k (CB, yellow) (right images). Scale bar: 30 μm. (**b**) qRT-PCR analysis of *Drd1, Drd2* and *Drd3* transcripts in cerebellar extracts from *Gfap-RiboTag* mice after HA-immunoprecipitation. All genes were normalized to *Tbp2a*. Data are presented as the fold change comparing the pellet fraction versus the input (n = 4-5 pooled samples of 2 mice / pool). Data were analyzed by twosided *t* test. \*\*\**p* < 0.001. (**c**) Representative immunoblots (left) and quantification (right) of GluA1 phosphorylation at S845 (pS845-GluA1) in the cerebellum of C57/Bl6 mice pretreated with the D1R/D5R antagonist, SCH23390 (0.1 mg/kg), 30 min prior saline or d-amphetamine (10 mg/kg) administration. Phosphorylated forms of GluA1 were normalized to unphosphorylated GluA1. β-actin was used as housekeeping protein. Data are expressed as percentage of saline group (n = 5-6 mice / group). Data are presented as means ± SEM and analyzed by one-way ANOVA followed by Tukey post-hoc comparisons test. \*\**p* < 0.01. (**d**) Representative immunoblots (left) and quantification (right) of GluA1 phosphorylation at S845 (pS845-GluA1) in the cerebellum of C57/Bl6 mice administered with saline, GBR12783 (15 mg/kg) and cocaine (20 mg/kg). Phosphorylated forms of GluA1 were normalized to unphosphorylated GluA1. β-actin was used as housekeeping protein. Data are expressed as percentage of saline group (n = 6 mice / group). Data are presented as means ± SEM and analyzed by one-way ANOVA followed by Tukey post-hoc comparisons test. PCl: Purkinje cell layer; g.l: granular layer; m.l: molecular layer. For detailed statistics see Supplementary Table 4.

**Figure 5.**
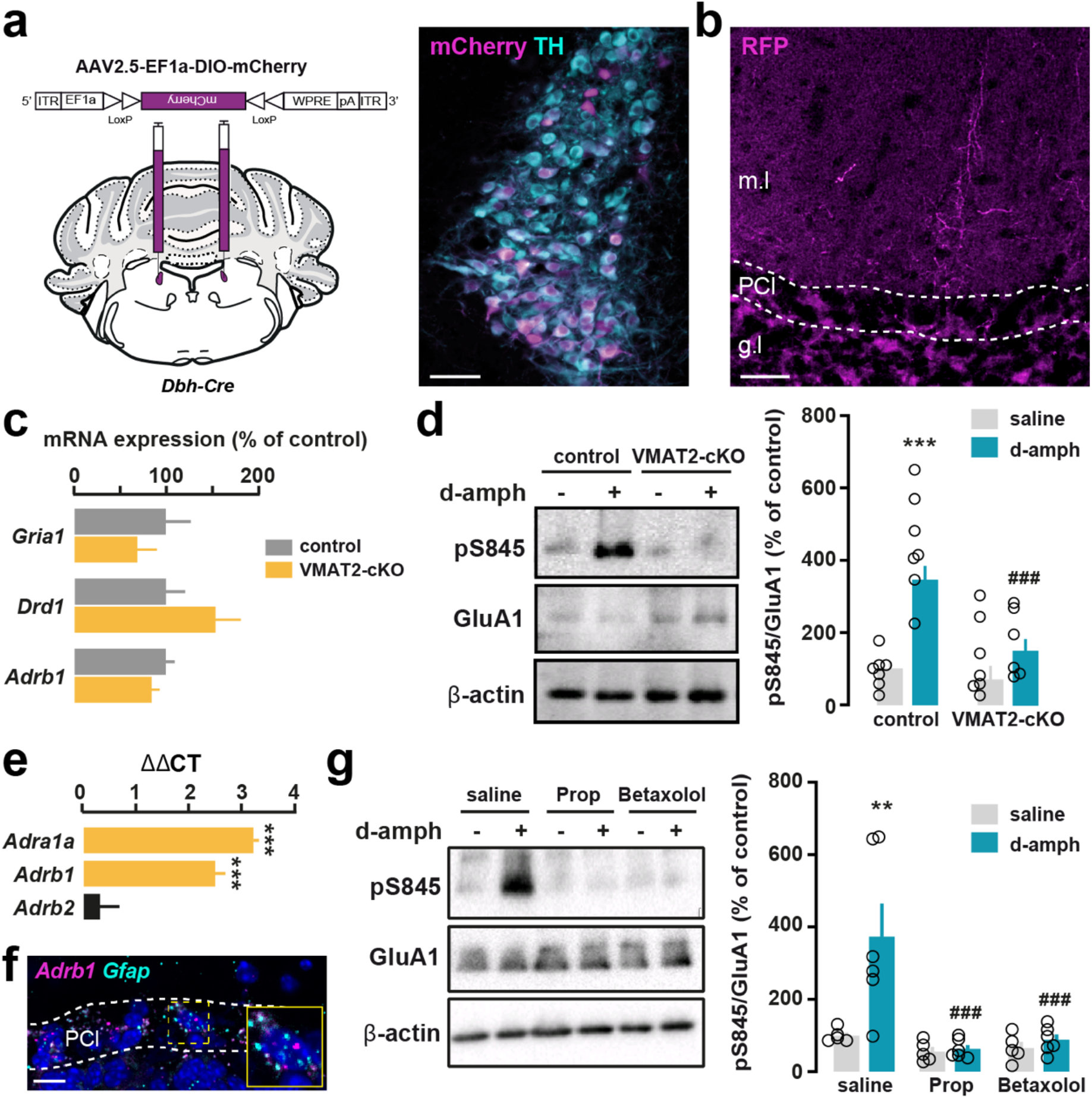
D-amphetamine-induced p845-GluA1 requires intact NE transmission and β1-AR activation. (**a**) Scheme of Cre-dependent AAV2.5-EF1a-DIO-mCherry injection in the LC of *Dbh-Cre* mice and representative images of transduced DBH neurons expressing mCherry (magenta) and TH (cyan). Scale bar: 50 μm. (**b**) RFP-expressing NE LC neuron axons positive fibers in the cerebellar cortex. Scale bar: 30 μm. (**c**) Expression of *Gria1, Drd1* and *Adrb1* transcripts in cerebellar extracts of control and VMAT2-cKO mice. All genes were normalized to *Tbp2a*. Data are expressed as % of control (n = 7 mice). Results are represented as mean ± SEM. Data were analyzed by two-sided *t* test. (**d**) Representative (right) and quantification (right) of GluA1 phosphorylation at S845 in the cerebellar extracts of control and VMAT2-cKO mice treated with saline or d-amphetamine (10 mg/kg). β-actin was used as housekeeping protein. Data is expressed as a percentage of control mice treated with saline (n = 6-7 mice / group). Results are represented as mean ± SEM and analyzed by two-way ANOVA followed by Bonferroni’s post-hoc comparisons. (**e**) qRT-PCR analysis of *Adrb1, Adrb2* and *Adra1a* transcripts in cerebellar extracts from *Gfap-RiboTag* mice after HA-immunoprecipitation. All genes were normalized to *Tbp2a*. Data are presented as the fold change comparing the pellet fraction versus the input (n = 5-6 pooled samples of 2 mice / pool). Data were analyzed by twosided *t* test. \*\*\**p* < 0.001. (**f**) Single molecule fluorescent *in situ* hybridization for *Adrb1* (magenta) and *Gfap* (cyan) mRNAs in the cerebellar cortex. The slide was counterstained with DAPI (blue). Note that *Adrb1* mRNAs are preferentially expressed in astrocytes in the PC layer most likely corresponding to somas of BG. Scale bar: 15 μm. (**g**) Representative (right) and quantification (right) of GluR1 phosphorylation at S845 in the cerebellar extracts of C57/Bl6 mice pretreated with the β1/2-AR antagonist propranolol (20 mg/kg) and β1-AR antagonist betaxolol (20 mg/kg), respectively, 30 min prior saline or d-amphetamine (10 mg/kg) administration. Phosphorylated forms of GluA1 were normalized to unphosphorylated GluA1. β-actin was used as housekeeping protein. Data is expressed as a percentage of control mice treated with saline (n = 5-6 mice / group). Results are represented as mean ± SEM and analyzed by one-way ANOVA followed by Tukey’s post-hoc comparisons. \*\**p* < 0.01 saline *vs*. d-amph, ### *p* < 0.001 sal/d-amph vs drug/d-amph. PCl: Purkinje cell layer; g.l: granular layer; m.l: molecular layer. For detailed statistics see Supplementary Table 4.

**Figure 6.**
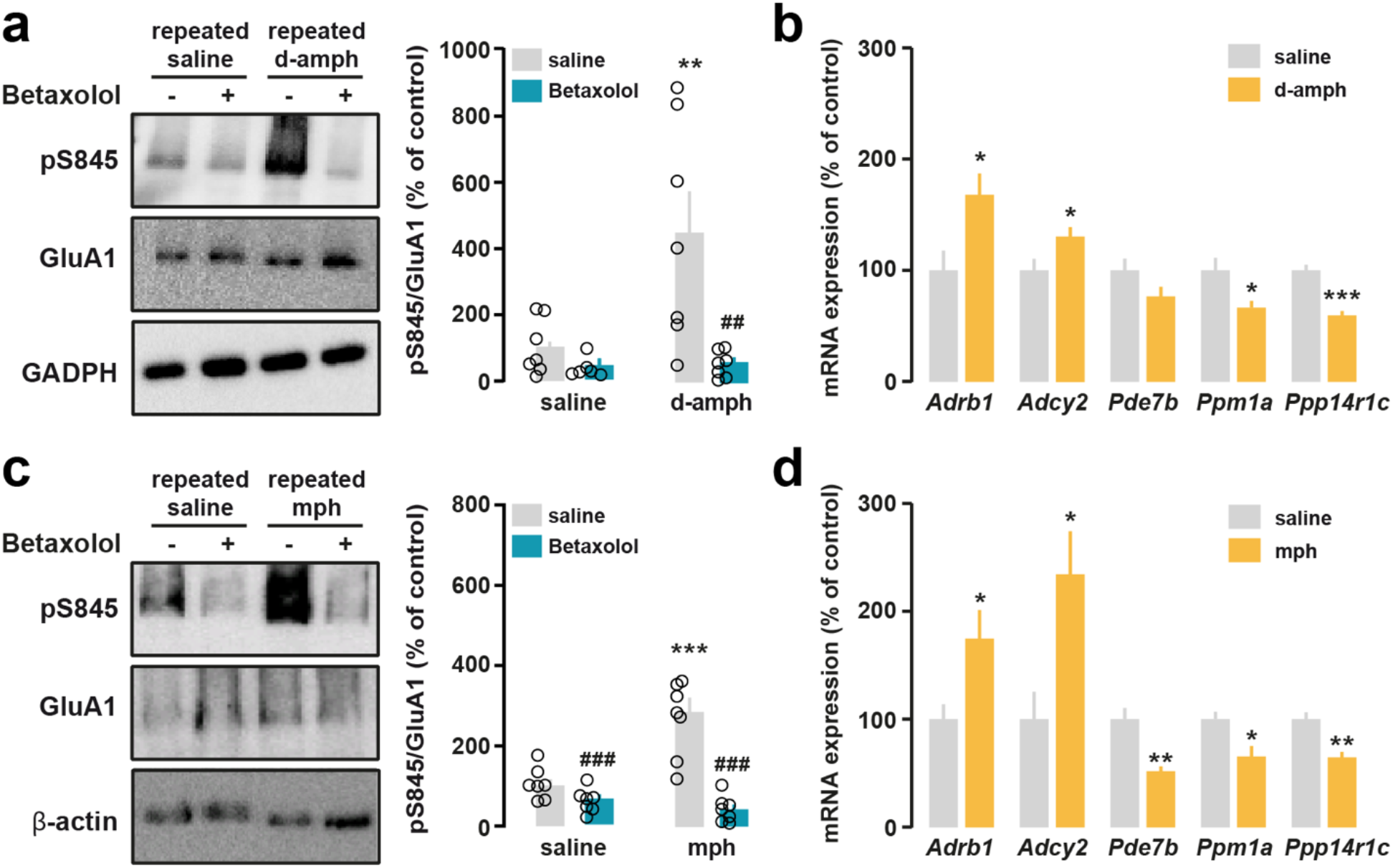
Effect of repeated exposure to d-amphetamine and methylphenidate on pS845-GluA1 in the cerebellum. (**a**) Representative immunoblots (left) and quantification (right) of GluA1 phosphorylation at S845 (pS845-GluA1) in the cerebellum of C57/Bl6 mice pretreated with betaxolol (20 mg/kg), 30 min prior each injection of saline or d-amphetamine (10 mg/kg, 1 inj/day during 5 days). Phosphorylated forms of GluA1 were normalized to unphosphorylated GluA1. GAPDH was used as housekeeping protein. Data are expressed as percentage of saline group (n = 6-7 mice/group). Data are presented as means ± SEM and analyzed by one-way ANOVA followed by Tukey post-hoc comparisons post. \*\**p* < 0.01 saline vs d-amph, ##*p* < 0.01 saline *vs*. betaxolol. (**b**) Comparison of the expression level of BG-enriched transcripts between mice repeatedly administered with saline or d-amphetamine. All genes analyzed by qRT-PCR were normalized to *Tbp2a*. Data are presented as mean ± SEM and analyzed by twosided *t* test. \**p* < 0.05, \*\*\**p* < 0.001. (**c**) Same analyses as in (**a**) performed in methylphenidate-treated mice. Data are expressed as percentage of saline group (n = 6-7 mice/group). Data are presented as means ± SEM and analyzed by one-way ANOVA followed by Tukey post-hoc comparisons post. \*\*\**p* < 0.001 saline *vs*. d-amph, ###*p* < 0.001 saline vs betaxolol. (**d**) Same analyses as in (**c**) performed in methylphenidate-treated mice. \**p* < 0.05, \*\**p* < 0.01. For detailed statistics see Supplementary Table 4.

### Tissue collection for polysome immunoprecipitation

Three weeks after the tamoxifen administration, male and female *Gfap-RiboTag* mice were killed by cervical dislocation and the heads were immersed in liquid nitrogen for 4 sec. The brains were then removed and sectioned on an aluminum block on ice and the whole cerebellum was rapidly isolated from the stem brain. Then, cerebellar samples were kept at −80°C until they have been used to performed polysome immunoprecipitation (IP).

### Polyribosome immunoprecipitation and RNA extraction

HA-tagged-ribosome immunoprecipitation in the cerebellum of *Gfap-RiboTag* mice was performed as it was previously described (Ceolin et al., 2017) using anti-HA antibody (5μl/sample; Biolegend; Cat#901502) and magnetic beads (Invitrogen, #100.04D). Total RNA contained in the pellet fraction was extracted from ribosome-mRNA complexes using RNeasy Microkit (Qiagen; Cat#74004) and from the input fraction using the RNeasy Minikit (Qiagen; Cat#74104) following manufacturer’s instructions. Quality and quantity of RNA samples were both assessed using the Nanodrop 1000 spectrophotometer. Between 5 and 9 biological replicates, each one composed of a pool of 2 mice, were used for qRT-PCR analysis (Figures 1e, 4b, 5e and Supplemental Figure 5). For the RNA extraction in Figures 5c, 6b and 6d and the Supplemental Figure 4 we used the RNAeasy Minikit (Qiagen; Cat#74104) following the manufacturer’s instruction.

### cDNA synthesis and quantitative real-time PCR

After RNA extraction from pellet and the input fractions of CC *Gfap-RiboTag* mice, synthesis of cDNA was performed as it was previously described by using the SuperScript VILO cDNA synthesis kit (Invitrogen) in one cycle program consisting of 10 min at 25°C, 60 min at 42°C, 5 min at 25°C and a final extension period of 5 min at 4°C. Resulting cDNA was used for quantitative real-time PCR (qRT-PCR), using SYBR Green PCR master mix on the LC480 Real-Time PCR System (Roche) and the primer sequences reported in Supplemental Table 3. Analysis was performed using LightCycler 480 Software (Roche). In Figures 1e, 3c, 4b, 5e and Supplemental Figure 4, the immunoprecipitated mRNA (pellet fraction) was compared to the input fraction. Results are presented as linearized Ct-values and normalized to the housekeeping *Tbp2a*. ΔΔCt method was used to give the fold change. Five to nine biological replicates were used in these experiments. In Figure 5c and Supplemental Figure 4, results are presented as % of change of VMAT2-cKO mice compared to control mice for each gene tested and normalized to the housekeeping gene *Tbp2a*. In Figure 6, results are presented as % of change of chronic d-amphetamine or methylphenidate versus chronic saline for each gene tested and normalized to the housekeeping gene *Tbp2a*.

### Single molecule fluorescent *in situ* hybridization

For the examination of targeted RNA within intact cells, in situ hybridization RNAscope^®^ technology was used following the protocol described by the supplier. Mice were decapitated and brains were frozen immediately on dry ice for 5 min and stored at −80°C. Brains were then sectioned at −17°C with a cryostat at 14 μm and mounted onto Superfrost Utra Plus slides (Thermo Scientific; Cat# J4800AMNZ). Coronal cerebellar sections were collected from bregma −5.80 mm and −6.80 mm. Probes for *Adrb1* (ACDBio; Mm-adrb1-C1Cat# 449761) and *Gfap* (ACDBio; Mm-gfap-C3 Cat#313211-C3) were used with the RNAscope Fluorescent Multiplex Kit (ACDBio; Cat# 320850) as described by the supplier. Slides were counterstained for DAPI and mounted with ProLong Diamond Antifade mountant (Invitrogen; Cat# P36961).

### Statistical analyses

GraphPad Prism v6.0 software was used for statistical analyses. For normally distributed data, Student’s t test was used. Multiple comparisons were performed by one-way or two-way ANOVA followed by Tukey’s post-hoc analyses. All data are presented as mean ± SEM, and statistical significance was accepted at 5% level. *p < 0.05, **p <0.01, ***p < 0.001. All the statistics are presented in the Supplementary Table 4.

## Results

### GluA1 is expressed in Bergmann glia cells in the adult mouse cerebellum

To confirm the preferential expression of GluA1 subunit of AMPA receptors in BGCs in the adult mouse cerebellum (Burnashev et al., 1992), we took advantage of the Ribotag methodology to isolate *Gria1* transcripts from BGCs. We first generated *Gfap-CreERT2-RiboTag* (*Gfap-RiboTag*) mice, which express the ribosomal protein Rpl22 tagged with hemagglutinin (HA) exclusively in astrocytes (**Figure 1**). Indeed, immunofluorescence analyses revealed that the vast majority of HA-positive cells had the typical morphology of BGCs with their cell bodies located in the PC layer and radial processes crossing the entire molecular layer. Moreover, these HA-positive cells co-localized with the astrocytic marker GFAP as well as with the glial glutamate/aspartate transporter GLAST-1 confirming their astrocytic nature (**Figure 1a-b**). In contrast, no co-localization was observed with neuronal (NeuN), Purkinje cells (PC) (Calbindin-D28k, DARPP-32) or microglial (Iba1) markers (**Figure 1c-d**). The specificity of *Gfap-RiboTag* mice was further validated by assessing the relative enrichment of astrocytic transcripts from mRNAs isolated after HA-immunoprecipitation on cerebellar extracts (**Supplemental Figure 1**). As expected, quantitative real time PCR (qRT-PCR) analyses revealed the enriched expression of several BGCs markers (*Gfap, S100b, Vim, Slc1a3, Cdc42ep4, Lgi4, Dao, Acsbg1*) (Koirala and Corfas, 2010) in the pellet fraction compared to the input fraction (**Figure 1e**). In contrast, transcripts which molecularly defined PC (*Pcp2, Calb1*), granule cells (*Neurod1*), Golgi cells (*Grm2*), unipolar brush cells (*Grp*), Lugaro cells (*Acan*), GABAergic interneurons (*Lypd6, Nos1ap*), microglia (*Aif1*), and oligodendrocytes (*Cnp*) were all depleted (**Figure 1e**). In this *Gfap-RiboTag* mouse model, qRT-PCR analysis revealed that *Gria1* transcripts were enriched after HA-immunoprecipitation (**Figure 1e**). The preferential expression of GluA1 in the molecular layer of the cerebellar cortex was confirmed by immunofluorescence (**Figure 1f**). Altogether, these results confirmed previous work revealing that GluA1 is preferentially expressed in BGCs in the adult mouse cerebellum (Petralia and Wenthold, 1992).

### D-amphetamine and methylphenidate increase GluA1 phosphorylation at serine 845 in the cerebellum

Acute administration of psychostimulant drugs, such cocaine or d-amphetamine triggers a rapid increase of GluA1 phosphorylation at serine 845 (pS845-GluA1) in the striatum and prefrontal cortex (Pascoli et al., 2005; Snyder et al., 2000; Valjent et al., 2005). To evaluate whether psychostimulants could produce similar effects in the cerebellum, C57/Bl6 mice were treated with d-amphetamine (10 mg/kg) and pS845-GluA1 was examined by Western blot in both the vermis and the lateral lobes of the cerebellum (**Figure 2**). Acute d-amphetamine administration, which produced hyperlocomotion and stereotypies within 15 min after injection (Pascoli et al., 2005), caused a rapid increase of pS845-GluA1 levels in the cerebellar vermis and lateral cerebellar hemispheres (**Figure 2a**). This effect was not accompanied by significant changes in the levels of total GluA1 detected with an antibody recognizing both the phosphorylated and unphosphorylated forms of GluA1 (**Figure 2a**). Moreover, subcellular fractionation of cerebellar lysates from mice injected with saline or d-amphetamine showed that pS845-GluA1 increases specifically in the in the membrane compartment (**Figure 2b**). In contrast, acute d-amphetamine failed to regulate GluA1 phosphorylation at serine 831 (pS831-GluA1) (**Figure 2a-b**). Similarly, an increase of pS845-GluA1 (p = 0.055) was observed in mice administered with methylphenidate (15 mg/kg), a psychostimulant drug commonly used for the treatment of ADHD (**Figure 2c**) (Wilens et al., 2002). Interestingly, a significant decrease of pS831-GluA1 was observed in the membrane fraction of methylphenidate-treated mice (**Figure 2c**). Altogether, these results indicate that acute d-amphetamine or methylphenidate administration selectively enhances GluA1 phosphorylation at serine 845 which most likely occurred in BGCs.

### Regulation of pS845-GluA1 in BGCs in response to d-amphetamine is independent from cerebellar dopamine signaling

D-amphetamine increases the extracellular concentration of dopamine (DA) availability in various brain areas (Kuczenski et al., 1997). We therefore conducted a series of experiments to determine whether DA signaling participates to the regulation of pS845-GluA1 by d-amphetamine in the cerebellum. Immunofluorescence analyses revealed the presence of a dense lattice of TH-positive fibers and a strong VMAT2 immunoreactivity in the molecular layer suggesting that DA could be potentially released in the cerebellar cortex (**Supplemental Figure 2a-b**). We first asked whether the VTA/SN constitute a source of dopaminergic fibers in the cerebellar cortex. To do so, midbrain DA axons were anterogradely labeled by injecting a Cre-dependent virus (AAV8-hSyn-FLEX-ChrimsonR-tdTomato) in the VTA/SN of *Slc6a3-Cre* mice, expressing the Cre recombinase under the promotor of the dopamine transporter (DAT) (**Figure 3a**). Viral transduction of midbrain DAT-positive neurons resulted in sparse axonal labeling in the PC and molecular layers detected using an antibody directed against the red fluorescent protein (RFP) (**Figure 3b and Supplemental Figure 3**). In the molecular layer RFP-positive fibers are oriented in the same axis of PC being in close proximity of BGCs identified using anti-GFAP antibody (**Figure 3b**). These results indicate that midbrain constitutes a potential source of dopaminergic input for the cerebellum.

In various brain regions, astrocytes express, although at low levels, transcripts encoding proteins involved DA metabolism (Juorio et al., 1993; Li et al., 1992). We therefore evaluated the expression of transcripts encoding enzymes involved in DA biosynthesis (*Ddc, Th*) as well as the dopamine transporter DAT (*Slc6a3*) among mRNAs isolated after HA-immunoprecipitation in cerebellar extracts of *Gfap-RiboTag* mice (**Figure 3c**). Our qRT-PCR analysis revealed that the expression of *Slc6a3* was enriched in cerebellar GFAP-positive cells (**Figure 3c**). In contrast, no changes were observed for *Th* and *Ddc* transcripts, which encode for the tyrosine hydroxylase and the aromatic l-amino acid decarboxylase, respectively (**Figure 3c**). The presence of DAT in BGCs was further supported by the analysis of YFP expression in the cerebellar cortex of *DAT-YFP* mice (**Figure 3d**). Thus, sparse cerebellar YFP-expressing cells, which displayed typical BGCs morphology, co-expressed the astrocytic marker GFAP but not the GABAergic marker PV (**Figure 3d**). These findings indicate that DAT is expressed by a fraction of BGCs raising the possibility that BGCs reuptake DA in the cerebellum. Altogether, our results identify at least two potential sites through which DA could be released and possibly participate to the regulation of pS845-GluA1.

In the striatum, d-amphetamine-induced pS845-GluA1 depends on cAMP-dependent protein kinase (PKA) activation downstream the stimulation of dopamine D1 receptors (D1R) (Snyder et al., 2000; Valjent et al., 2005). Despite early evidence suggesting the presence of D1R in the molecular layer of the cerebellum (Camps et al., 1990; Savasta et al., 1986), the cellular identity of D1R-expressing cells in the cerebellar cortex remains largely unknown. To address this issue, we analyzed the distribution of GFP-positive cells in the cerebellar cortex of *D1-eGFP* mice (**Figure 4**). In all the lobules, a strong GFP labeling was found in the molecular layer of the cerebellar cortex (**Figure 4a**). Detailed analysis revealed that GFP-positive cells expressed GFAP, a marker of astrocytes, presumably corresponding to BGCs, but not the PC marker CB (**Figure 4a**). The presence of D1R in BGCs was further supported by the enrichment of *Drd1* transcripts among the mRNAs isolated following HA-immunoprecipitation in cerebellar extracts of *Gfap-RiboTag* mice (**Figure 4b**). In contrast, no differences were observed for *Drd2* and *Drd3* transcripts (**Figure 4b**). Together, these results indicate that D1R are expressed in BGCs.

To assess whether D1R participates to the regulation of GluA1 phosphorylation induced by d-amphetamine in BGCs, C57/Bl6 mice were administered with SCH23390 (0.1 mg/kg), a D1R/D5R-selective antagonist, 30 min prior the injection of d-amphetamine or saline. As shown in Figure 4, blockade of D1R/D5R had no effect on the basal or increased GluA1 phosphorylation at S845 induced by d-amphetamine measured in membrane fractions (**Figure 4c**). These results suggest that DA may not be necessary to enhance GluA1 phosphorylation in BGCs in response to d-amphetamine. We therefore evaluated whether increasing the extracellular concentration of DA was sufficient to trigger pS845-GluA1 in the cerebellum. To do so, we measured the effect of cocaine (20 mg/kg) and GBR12783 (15 mg/kg), a selective DA reuptake inhibitor, on the phosphorylation of GluA1 at S845. Although both drugs enhanced locomotor activity 15 min after administration (Valjent et al., 2010), they failed to increase pS845-GluA1 in BG (**Figure 4d**). Similar results were obtained following the acute administration of the D1R/D5R agonist, SKF81297 (data not shown). Altogether, our findings indicate that d-amphetamine-induced GluA1 phosphorylation at S845 in BGCs does not involve DA transmission.

### D-amphetamine-induced increase of pS845-GluA1 requires noradrenergic transmission

In addition to its ability to release DA, d-amphetamine is also a potent releaser of norepinephrine (Kuczenski and Segal, 1997). We therefore examined whether the cerebellar cortex received noradrenergic-projecting neurons from the locus coeruleus (LC), the major noradrenergic nucleus of the brain (Saigal et al., 1980; Saint-Mleux et al., 2004). To selectively label hindbrain NE axons, we injected a Cre-dependent virus (AAV2.5-EF1a-DIO-mCherry) in the LC of mice expressing the Cre-recombinase under the dopamine β-hydroxylase promoter (*Dbh-*Cre mice) (**Figure 5a**). As shown in Figure 5, RFP-positive fibers were detected in the PC and molecular layers indicating that NE neurons from the LC constitute a source of noradrenergic input for the cerebellum (**Figure 5b**).

We therefore investigated whether the noradrenergic transmission was involved in the increased pS845-GluA1 induced by d-amphetamine administration. To address this issue, we used conditional *Slc18a2* knock-out mice (VMAT2-cKO) previously generated by crossing the *Dbh*-Cre mouse line with *Slc18a2^loxP/loxP^* mice (Isingrini et al., 2016). As revealed by qRT-PCR analysis performed on cerebellar extracts, deletion of *Slc18a2* in DBH neurons neither altered level of transcripts encoding receptors (*Drd1, Drd2, Drd3, Adrb1, Adrb2, Adra1a*) transporters (*Slc6a3*), and enzymes involved in catecholamine biosynthesis (*Ddc, Th*) and degradation (*Comt, Maoa, Maob*), nor changed the expression of *Gria1* transcripts (**Figure 5c and Supplemental Figure 4**). In addition, no changes were detected in the expression of transcripts enriched in glial cells including BGCs (*Gfap, Itgam, Aif1, Apq4, Kcnj10*) and PC (*Gria2\* We then measured d-amphetamine-induced pS845-GluA1 in VMAT2-cKO. While cerebellar GluA1 expression was unaffected by the deletion of *Slc18a2* in the DBH neurons, the increase of pS845-GluA1 observed following d-amphetamine administration was totally abolished in VMAT2-cKO (**Figure 5d**). These results indicate that noradrenergic transmission is necessary for the regulation of GluA1 phosphorylation by d-amphetamine in BGCs.

We next investigated whether d-amphetamine-induced PKA-dependent GluA1 phosphorylation in BGCs required β-Adrenergic receptors (β-AR) which, upon agonist activation stimulate the cAMP/PKA pathway (Gelinas et al., 2008; Maguire et al., 1977). Using mRNAs isolated after HA-immunoprecipitation in cerebellar extracts of *Gfap-RiboTag* mice, we determined the relative abundance of *Adrb1* and *Adrb2* transcripts encoding β1-AR and β2-AR, respectively (**Figure 5e**). Our analysis revealed that *Adrb1*, but not *Adrb2*, transcripts were significantly enriched in cerebellar GFAP-positive cells (**Figure 5e**). Of note, *Adra1a* transcripts encoding a1a-AR, which are highly expressed in BGCs (Doyle et al., 2008), were also detected after HA-immunoprecipitation (**Figure 5e**). Single molecule fluorescent *in situ* hybridization analysis further confirmed the preferential expression of *Adrb1* mRNAs in BGCs, identified here by *Gfap* transcripts in the PC layer (**Figure 5f**). To evaluate the contribution β1-AR in the regulation of GluA1 phosphorylation triggered by d-amphetamine, C57/Bl6 mice were administered with either propranolol, a general β-AR antagonist (20 mg/kg; i.p), or betaxolol, a selective β1-AR antagonist (20 mg/kg; i.p) 30 min prior the injection of d-amphetamine or saline (**Figure 5g**). As revealed by Western blot analyses, d-amphetamine failed to increased GluA1 phosphorylation in presence of both antagonists (**Figure 5g**). Altogether, these findings indicate that d-amphetamine-induced pS845-GluA1 in BG requires noradrenergic transmission and β1-AR activation.

### Repeated exposure to d-amphetamine and methylphenidate enhances pS845-GluA1 in BGCs through β1-AR activation

Long-term exposure to d-amphetamine or methylphenidate induce a variety of neuronal changes that contribute to the development of long-lasting behavioral alterations (refs). We next investigated whether the ability of d-amphetamine to trigger pS845-GluA1 was preserved following repeated exposure and if so, whether it still relied on β1-AR activation. To address this issue, C57/Bl6 mice received saline or d-amphetamine (10 mg/kg) for 5 consecutive days. Before each administration, mice were pretreated with betaxolol (20 mg/kg) or its vehicle. The level of pS845-GluA1 was analyzed in cerebellar membrane fractions 15 min after the last injection of saline or d-amphetamine. Western blot analysis revealed that the ability of d-amphetamine to increase pS845-GluA1 in BGCs was preserved in mice repeatedly exposed to d-amphetamine (**Figure 6a**). This increased phosphorylation still relied on β1-AR activation since pS845-GluA1 was totally prevented in mice pretreated with betaxolol (**Figure 6a**).

We next examined whether repeated exposure to d-amphetamine induced transcriptional alterations of genes encoding proteins involved in the regulation of cAMP/PKA pathway turnover. To address this question, we compared by qRT-PCR the expression level of several BGCs-enriched transcripts between mice repeatedly administered with saline or d-amphetamine (Figure 6b and Supplemental Figure 5). Our analysis revealed a significant increase of *Adrb1* and *Acdy2* transcripts encoding β1-AR and adenylate cyclase 2 in mice exposed 5 consecutive days to d-amphetamine (Figure 6b). Conversely, *Pde7b, Ppm1a* and *Ppplrl4c* mRNAs encoding phosphodiesterase 7B, protein phosphatase, Mg^2+^/Mn^2+^ dependent 1A and protein phosphatase 1 regulatory subunit 14C, respectively, were reduced upon this d-amphetamine-treatment regimen (Figure 6b). Similar results were obtained in mice repeatedly exposed to methylphenidate (Figure 6c-d). Altogether, these results indicate that β1-AR-dependent GluA1 phosphorylation in BGCs is preserved in mice repeatedly exposed to d-amphetamine or methylphenidate. Moreover, our findings identify BGCs transcriptional alterations of several regulators of the cAMP/PKA pathway, which account for the absence of pS845-GluA1 desensitization.

## Discussion

The present findings show that systemic administration of d-amphetamine and methylphenidate enhanced cAMP/PKA-regulated phosphorylation of GluA1 subunit of the AMPA receptors in BGCs. They also revealed that this regulation requires intact NE transmission and the activation of β1-AR. Finally, our study identified transcriptional alterations of several components of the cAMP/PKA pathway which may account for the maintenance of the stimulant medications ability to increase GluA1 phosphorylation following drug administration. These results suggest that BGCs may represent a key component in the efficacy of these drugs to alleviate ADHD symptoms.

GluA1 phosphorylation is an important process regulating AMPA receptors functions in response to a variety of stimuli (Roche et al., 1994). The regulation of GluA1 phosphorylation in response to psychostimulants is certainly one of the best characterized. Here we demonstrate a rapid and selective increase of GluA1 phosphorylation at S845 in the cerebellum following d-amphetamine and methylphenidate administration. Our results extend previous observations identifying similar regulatory mechanisms in other brain regions by a wide range of psychostimulants (Choi et al., 2011; Li et al., 2011; Mao et al., 2013; Pascoli et al., 2005; Snyder et al., 2000; Valjent et al., 2005; Xue et al., 2014). Importantly, our data suggest that each psychostimulant generate specific patterns of GluA1 phosphorylation, as it is the case for ERK pathway (Valjent et al., 2004 57). Thus, increased pS845-GluA1 in the cerebellum was not induced by cocaine but instead only observed following the administration of stimulant medications. This contrasts with the striatum, where enhanced GluA1 phosphorylation occurs in response to all psychostimulants (Ferrario et al., 2011; Mao et al., 2013; Snyder et al., 2000; Valjent et al., 2005). The ability of cocaine vs stimulant medications to recruit distinct monoaminergic circuits certainly account for the specificity of GluA1 phosphorylation patterns induced by these drugs. Thus, in densely DA-innervated regions such as the striatum, psychostimulants-induced pS845-GluA1 is mediated primarily through D1R activation (Mao et al., 2013; Snyder et al., 2000; Valjent et al., 2005). In contrast, our study reveals that in the cerebellum, in which both DA and NE extracellular concentrations increased following d-amphetamine and methylphenidate administration (Goldstein and Macmillan, 1993; Krobert et al., 1994; Quansah et al., 2018), GluA1 phosphorylation relies exclusively on intact NE transmission and β1-AR activation as it does in the prefrontal cortex (Pascoli et al., 2005; Xue et al., 2014). Interestingly, hippocampal GluA1 phosphorylation, which is enhanced by d-amphetamine (Mao et al., 2015), is also strongly regulated by NE and β1-AR agonist (Hu et al., 2007; Tenorio et al., 2010) suggesting that the contribution of the noradrenergic system in the regulation of pS845-GluA1 by stimulant medications is certainly not restricted to the cerebellum and the prefrontal cortex.

Psychostimulant-induced pS845-GluA1 in the striatum and the prefrontal cortex occur in D1R- and β1-AR-containing neurons (Pascoli et al., 2005; Snyder et al., 2000; Valjent et al., 2005). Our data clearly indicate that the ability of d-amphetamine or methylphenidate to enhance GluA1 phosphorylation is not specific to neurons but can also occur in astrocytes. In contrast to other brain regions, the expression of GluA1 subunit in the cerebellum is preferentially restricted to BGCs (Burnashev et al., 1992; Douyard et al., 2007; Matsui et al., 2005). In these glial cells GluA1-GluA4 containing AMPA receptors play an important role in regulating the astrocytic coverage of PC glutamatergic synapses (De Zeeuw and Hoogland, 2015; Ikai et al., 1994; Saab et al., 2012). Whether GluA1 phosphorylation at S845 in BGCs potentiates AMPA currents (Roche et al., 1996) and/or regulates surface trafficking of GluA1 (Serulle et al., 2007) as in neurons, remains to be established. Moreover, future studies will be necessary to determine whether d-amphetamine-induced pS845-GluA1 in BGCs is causally linked to the long-lasting β-AR-dependent reduction of PC firing rate induced by acute and repeated d-amphetamine exposure (Freedman and Marwaha, 1980; Sorensen et al., 1985; Sorensen et al., 1982).

Recent evidence indicates that depending on the vigilance states, astrocytes of the auditory cortex integrate NE activity through distinct signaling pathways. Thus, while transient NE release is accompanied with large cytosolic astrocytic Ca^2+^ elevations, sustained activity of noradrenergic neurons leads to a gradual increase of cAMP (Oe et al., 2020). Our results strongly suggest that similar regulations occur in BGCs since β1-AR-mediated GluA1 phosphorylation induced by stimulant medications relies on sustained activity on LC NE neurons projecting to the cerebellum. Despite the lack of regulation of GluA1 phosphorylation at the calcium-dependent site S831, BGCs Ca^2+^ dynamics might also be modulated in response to psychostimulants. Thus, α1-AR-dependent BGCs Ca^2+^ elevation has been reported during locomotion (Paukert et al., 2014), a behavioral response highly enhanced by d-amphetamine and methylphenidate exposure (Valjent et al., 2010). Moreover, acute d-amphetamine administration increases Ca^2+^ signaling in astrocytes in the nucleus accumbens though astrocytic D1R activation. Such regulatory mechanism is functionally important since it contributes to the modulation of both excitatory synaptic transmission and acute hyperlocomotor effects of d-amphetamine (Corkrum et al., 2020). Although D1Rs expressed in BGCs do not participate to the regulation of GluA1 phosphorylation, their potential role in the regulation of BGCs Ca^2+^ signaling will require further investigation.

Structural and functional abnormalities in the cerebellum were amongst the first reported in patients diagnosed for ADHD. Thus, the severity of ADHD symptoms correlates to the extent of cerebellar volume reduction, which can be compensated by methylphenidate treatment (Bledsoe et al., 2009; Castellanos et al., 2002; Ivanov et al., 2014; Rubia et al., 2009). Interestingly, increased metabolic activity could be one of the mechanisms by which methylphenidate normalizes cerebellar dysfunction observed in ADHD (Volkow et al., 1998; Volkow et al., 1997). Supporting this hypothesis, methylphenidate has been shown to increases glutamate uptake in BGCs contributing to normalize impaired metabolic homeostasis reported in ADHD (Guillem et al., 2015). Future studies will determine whether disruption of cerebellar glutamate homeostasis may contribute to development of substance use disorders associated with the non-medical use of stimulant medications.

## Abbreviations

BGCs: Bergmann Glia Cells
PC: Purkinje cells
CC: Cerebellar Cortex
DA: Dopamine
NE: Noradrenaline
D1R: Dopamine D1 receptor
β-AR: Beta-adrenergic receptors
β1-AR: Beta-1-adrenergic receptors
AC: Adenylyl cylase
PKA: Protein Kinase A
PKC: Protein Kinase C
AMPA: α-amino-3-hydroxy-5-methyl-4-isoxazolepropionic acid
GluA1: AMPA subunit A1
DAT: Dopamine transporter
TH: Tyrosine hydroxylase
VMAT2: Vesicular monoamine transporter 2
VTA: Ventral Tegmental Area
SN: Substantia nigra
LC: Locus Coeruleus
D-amph: D-amphetamine
Mph: Methylphenidate
ADHD: attention-deficit/hyperactivity disorder
HA: hemagglutinin

## Acknowledgements

We thank the iExplore, MRI and PVM Platforms of the IGF for their involvement in the maintenance and breeding of the colonies. This work was supported by Inserm, Fondation pour la Recherche Médicale (DEQ20160334919), La Marató de TV3 Fundació, ANR EPITRACES, ANR DOPAFEAR (EV), NARSAD Young Investigator Grant from the Brain and Behavior Research Foundation (EP) and CIHR project grant 201803PJT (BG). Laura Cutando is supported by the post-doctoral Labex EpiGenMed fellowship («Investissements d’avenir» ANR-10-LABX-12-01), EP was a recipient of Marie Curie Intra-European Fellowship IEF327648 and is currently a recipient of a Beatriu de Pinós fellowship (# 2017BP00132) from the University and Research Grants Management Agency (Government of Catalonia, Spain). Laia Castell is supported by the PhD Labex EpiGenMed fellowship («Investissements d’avenir» ANR-10-LABX-12-01). P.B and AKE are Research Associate and Research Director of the FRS-FNRS respectively and both are supported by FNRS, AKE is also supported by Foundation Simone and Pierre Clerdent.

## Author contributions

Laura Cutando, E.P and E.V conceived, designed and led the project. Laura Cutando and E.P performed brain dissections. Laura Cutando and E.P performed polysome IP and qRT-PCR experiments. Laura Cutando and F.B performed Western blot analyses. Laura Cutando, E.P and P.T performed immunofluorescence experiments. Laura Cutando and Laia Castell performed *in situ* hybridization analysis. M.G and G.D provided *Gfap-CreERT2* mice and reagents. P.B and A.K performed stereotaxic injections in *Dbh-Cre* mice. V.P and C.L provided *Slc6a3-Cre* injected mice. E.I. and B.G generated and provided VMAT2-cKO mice. E.V supervised the project. Laura Cutando and E.V wrote the manuscript with input from all authors. The authors declare no conflicts of interest.

## Supplemental Figure Legends

**Supplemental Figure 1.**
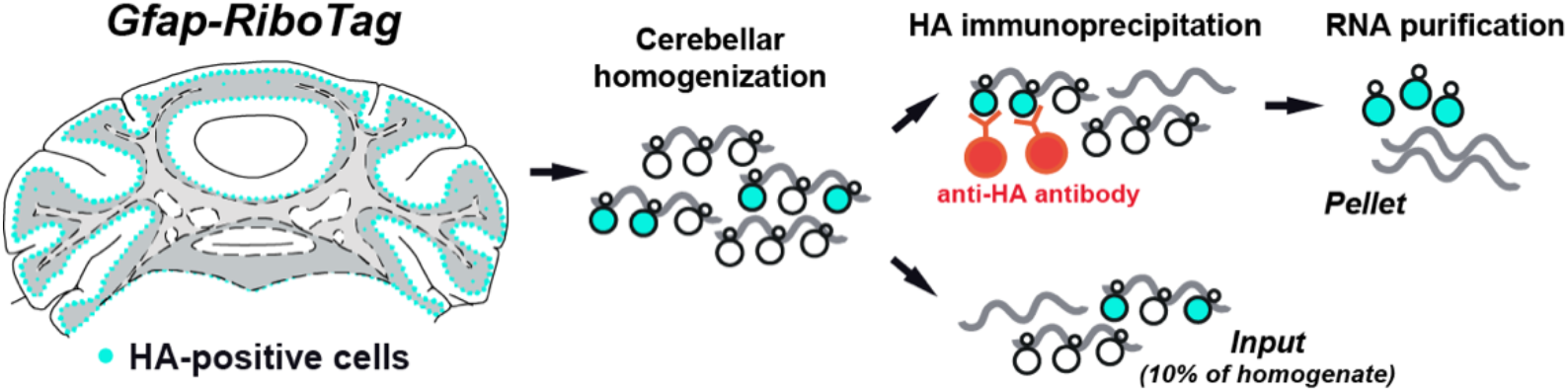
Ribotag methodology. Drawing illustrating the Ribotag methodology in the *Gfap-RiboTag* mice.

**Supplemental Figure 2.**
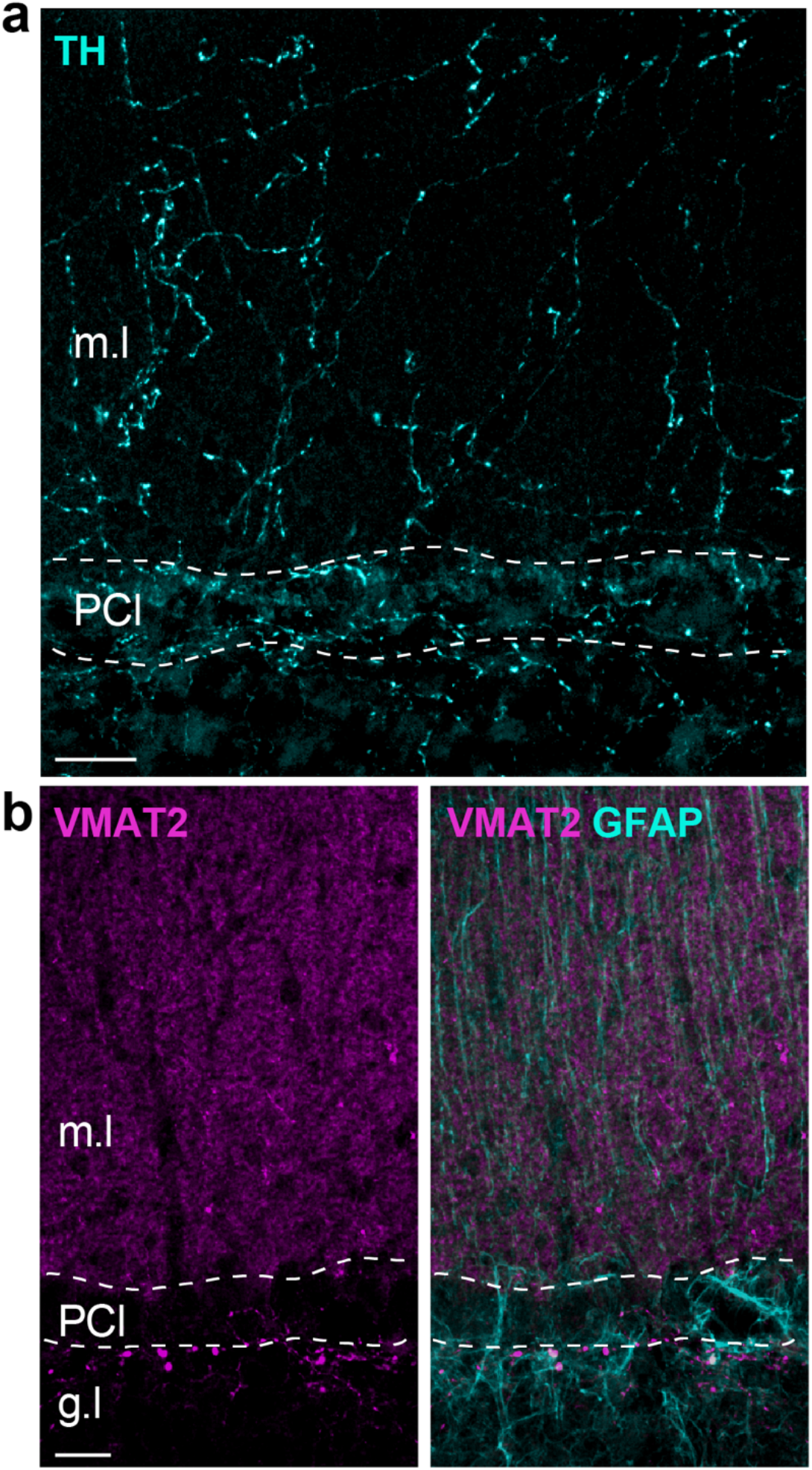
Distribution of TH- and VMAT2-positive fibers in the cerebellum. (**a**) Immunofluorescence for tyrosine hydroxylase (TH, cyan) in the cerebellar cortex. Note the lattice of TH-positive fibers present in both Purkinje (PC) and molecular layers (m.l). Scale bar: 30 μm. **(b**) Double immunofluorescence for VMAT2 (magenta) and GFAP (cyan). Note the dense VMAT2 immunoreactivity in the molecular layer. Scale bar: 15 μm. PCl: Purkinje cell layer; g.l: granular layer; m.l: molecular layer.

**Supplemental Figure 3.**
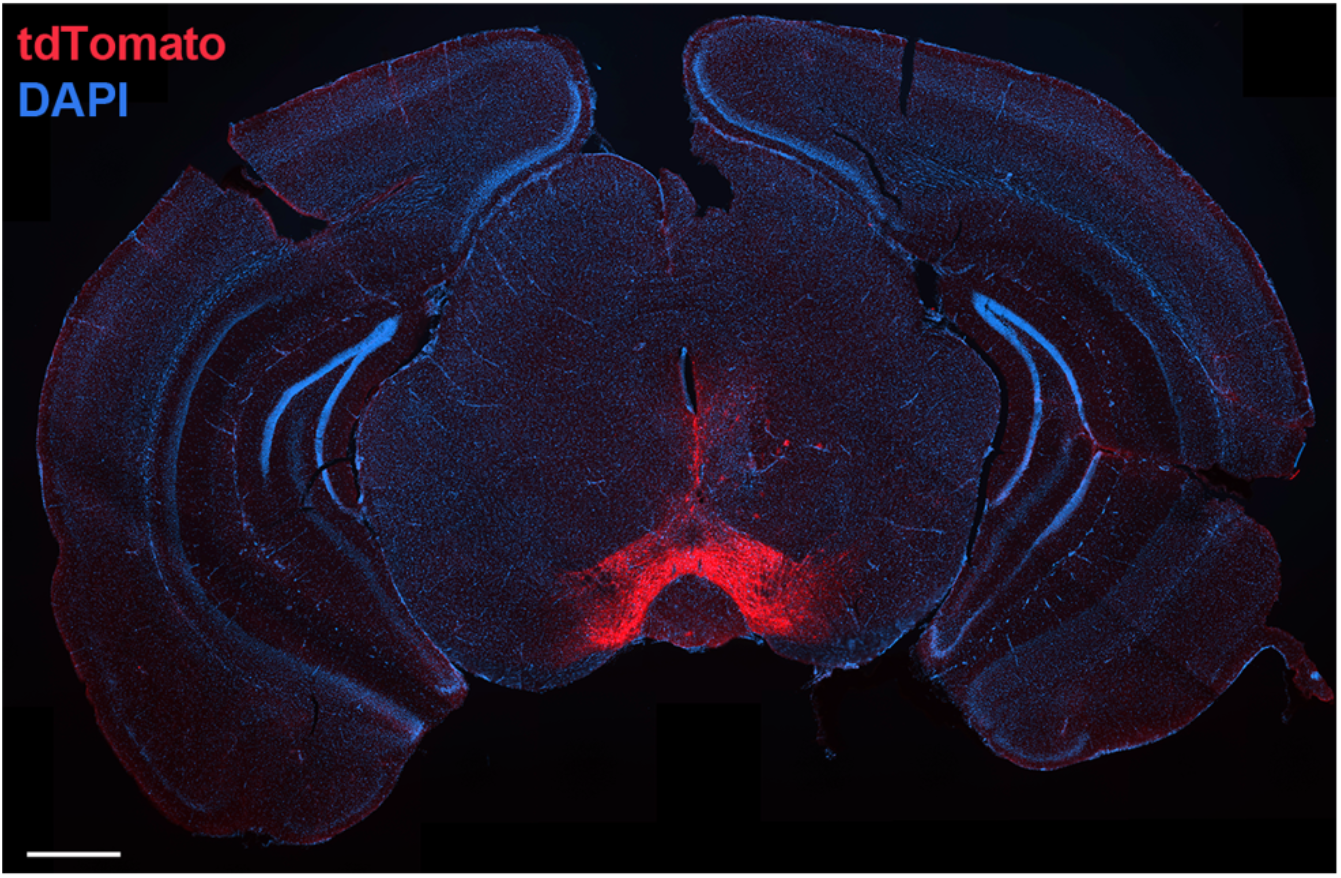
Injection of the amber light-drivable channelrhodospinChrimson in the VTA of *Slc6a3-Cre* mice. Representative images of transduced DAT neurons expressing tdTomato (red). Scale bar: 500 μm.

**Supplemental Figure 4.**
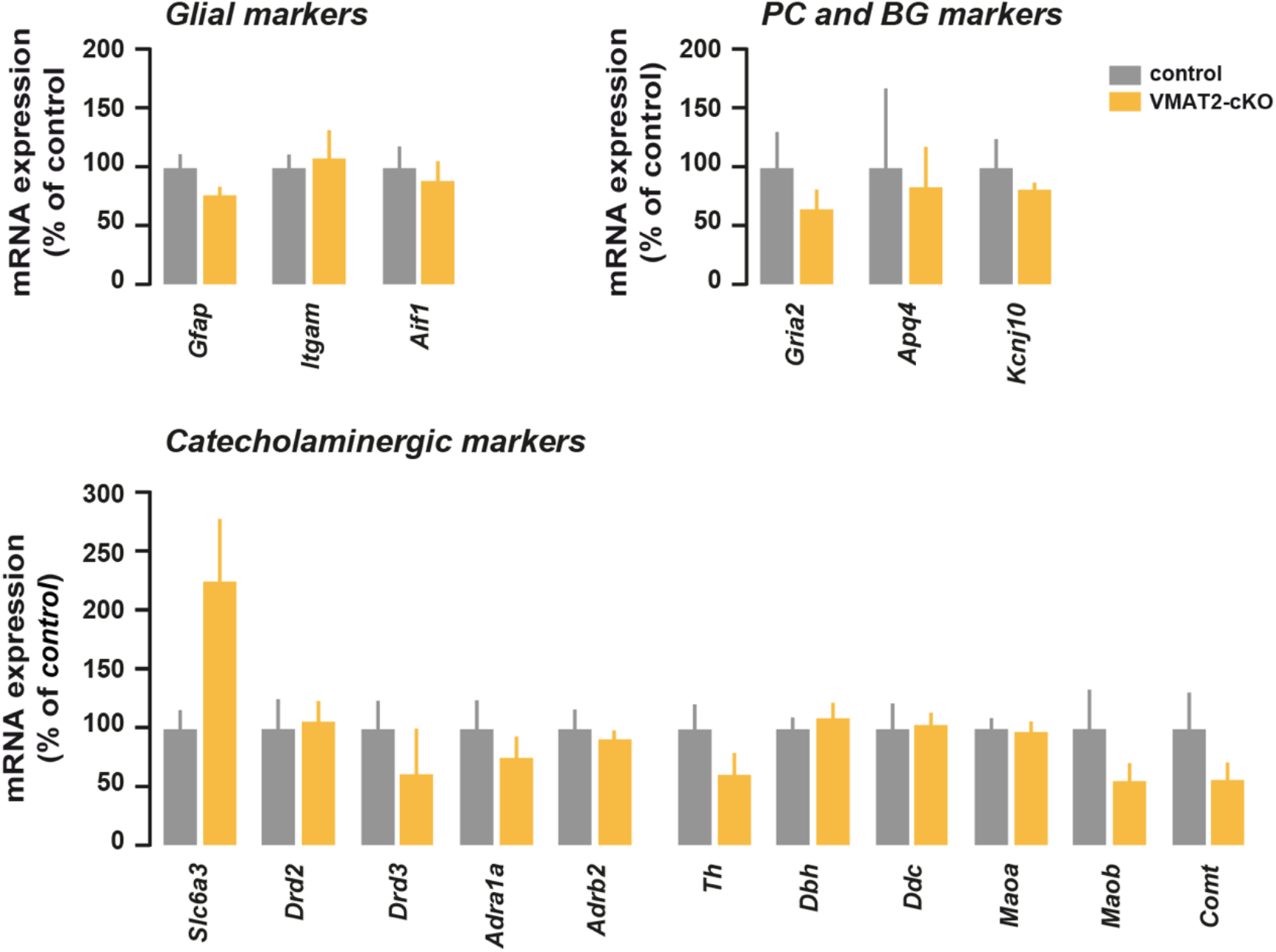
Transcriptional analysis in the cerebellum of VMAT2-cKO mice. Analysis of transcripts encoding glial cells (*Gfap, Itgam, Aif1*) including BGCs (*Apq4, Kcnj10*) and PC (*Gria2*) markers as well as transcripts encoding receptors (*Drd2, Drd3, Adrb2, Adra1a*) transporters (*Slc6a3*), and enzymes involved in catecholamine biosynthesis (*Ddc, Th*) and degradation (*Comt, Maoa, Maob*) in control and VMAT2-cKO mice (n = 6-7 mice / genotype). All genes were normalized to *Tbp2a*. Data are presented as mean ± SEM and analyzed by twosided *t* test. For detailed statistics see Supplementary Table 4.

**Supplemental Figure 5.**
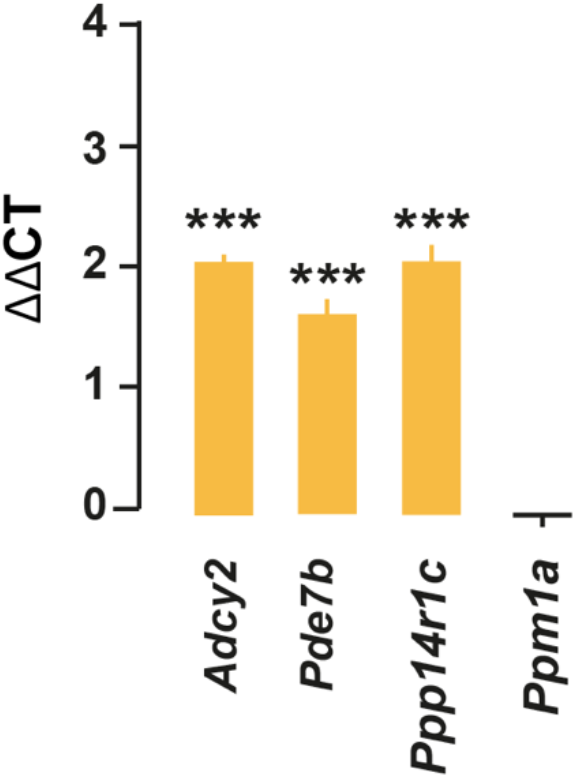
Transcripts encoding proteins involved in cAMP/PKA turnover enriched in BG. qRT-PCR analysis of *Adcy2, Pde7b, Ppp1r14c* and *Ppm1a* transcripts in cerebellar extracts from *Gfap-RiboTag* mice after HA-immunoprecipitation. All genes were normalized to *Tbp2a*. Data are presented as the fold change comparing the pellet fraction versus the input (n = 8-9 pooled samples of 2 mice / pool). Data were analyzed by two-sided *t* test. \*\*\**p* < 0.001. For detailed statistics see Supplementary Table 4.

**Supplementary Table 1.** Mouse lines used in the study.

Mouse lines used in the study
| Article's nomenclature (Mouse line) | Mouse strain | Citation |
| --- | --- | --- |
| ***D1R-eGFP*** (*Drd1a-eGFP*) | Tg(Drd1-EGFP)X60Gsat/Mmmh | (Gong et al., 2003) |
| ***Dbh-Cre*** (*Dbh-Cre*) | Tg(Dbh-cre)KH212Gsat/Mmucd | (Gong et al., 2003) |
| ***VMAT2-cKO***<br>(*Dbh-Cre:Slc18a2-LoxP/LoxP*) |  | (Isingrini et al., 2016) |
| ***Slc6a3-Cre*** (*Slc6a3-Cre*) | Tg(Slc6a3-icre)1Fto | (Turiault et al., 2007) |
| ***DAT-YFP***<br>(*Slc6a3-Cre:ROSA26-eYFP*) |  | (Madisen et al., 2010) |
| ***Gfap-RiboTag***<br>(*Gfap- CreERT2:RiboTag*) |  | (Mongrédien et al., 2019) |

**Supplementary Table 2.** List of Primary Antibodies.

List of Primary Antibodies
| Antigen | Species | Dilution | Supplier/Catalog no./References |
| --- | --- | --- | --- |
| HA | Mouse | 1:1000 (IF) | Biologend (#901502) |
| HA | Rabbit | 1:1000 (IF) | Rockland (#600-401-384) |
| DARPP-32 | Rabbit | 1:1000 (IF) | Cell Signaling Technology (#2306) |
| GFP | Chicken | 1:1000 (IF) | Life Technologies (#A10262) |
| RFP | Rabbit | 1:1000 (IF) | MBL (#PM005) |
| RFP | Mouse | 1:1000 (IF) | MBL (#M155-3) |
| Parvalbumin | Mouse | 1:1000 (IF) | Millipore (MAB1572) |
| TH | Rabbit | 1:1000 (IF) | Millipore (#AB152) |
| TH | Chicken | 1:1000 (IF) | Aves Labs |
| GFAP | Chicken | 1:1500 (IF) | Abcam (#ab4674) |
| GFAP | Rabbit | 1:1500 (IF) | Dako (#N1506) |
| VMAT2 | Rabbit | 1:1000 (IF) | Synaptic System (#138302) |
| Iba1 | Rabbit | 1:1000 (IF) | Wako (#019-19741) |
| Calbindin-D28k | Mouse | 1:1000 (IF) | Swant (#300) |
| Calbindin-D28k | Rabbit | 1:1000 (IF) | Swant (#CB38) |
| EAAT1 (GLAST-1) | Rabbit | 1:500 (IF) | Abcam (#ab416) |
| NeuN | Mouse | 1:1000 (IF) | Millipore (#MAB377) |
| GluA1 | Rabbit | 1:1000 (IF); 1:1000 (WB) | Millipore (#AB1504) |
| pS845-GluA1 | Rabbit | 1:1000 (WB) | Millipore (#04-1073) |
| pS831-GluA1 | Rabbit | 1:1000 (WB) | Millipore (#01-823) |
| β-actin | Mouse | 1:40000 (WB) | Abcam (#AB6276) |
| GAPDH | Mouse | 1:40000 (WB) | Santa Cruz (#SC-32233) |

**Supplementary Table 3.**
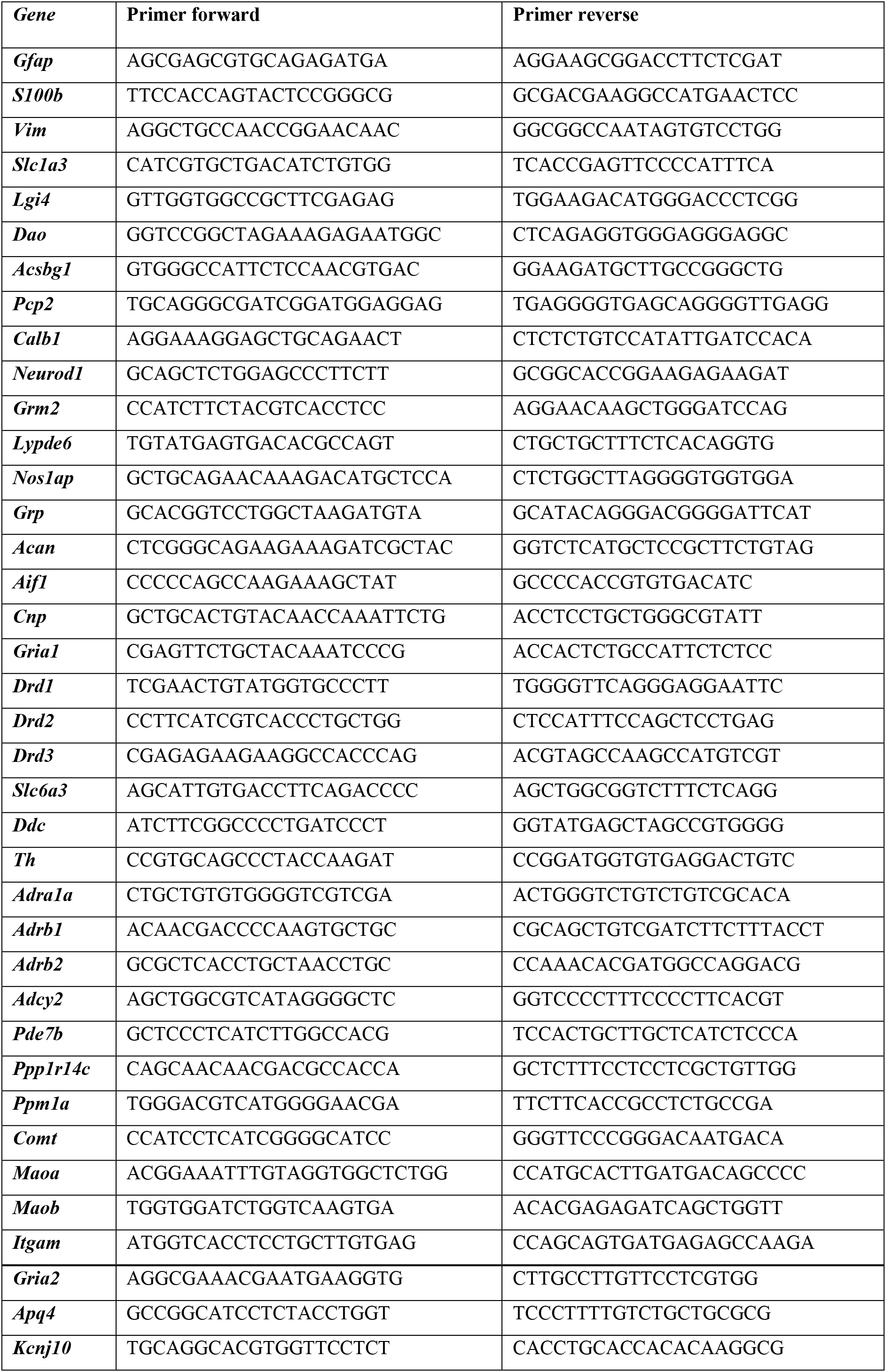
Sequences of PCR primers.

**Supplementary Table 4:** Statistical analysis.

| Figures | Groups (n number of mice) | Statistical Analysis |
| --- | --- | --- |
| 1e | *Transcripts* (input, pellet)<br>*Gfap* : (n = 5, n = 5)<br>*S100b* : (n = 5, n = 5)<br>*Vim* : (n = 6, n = 5)<br>*Slc1a3* : (n = 6, n = 5)<br>*Cdc42ep4* : (n = 6, n = 5)<br>*Lgi4* : (n = 6, n = 5)<br>*Dao* : (n = 6, n = 5)<br>*Acsbg1* : (n = 6, n = 5)<br>*Pcp2* : (n = 6, n = 5)<br>*Calb1* : (n = 6, n = 5)<br>*Neurod1* : (n = 6, n = 5)<br>*Grm2* : (n = 6, n = 5)<br>*Lypd6* : (n = 6, n = 5)<br>*Nos1ap* : (n = 6, n = 5)<br>*Grp* : (n = 6, n = 5)<br>*Acan* : (n = 6, n = 5)<br>*Aif1* : (n = 6, n = 5)<br>*Cnp* : (n = 6, n = 5)<br>*Gria1* : (n = 6, n = 5) | Unpaired t-test<br>$t_8 = 8.964$, $p < 0.0001$<br>$t_8 = 15.53$, $p < 0.0001$<br>$t_9 = 2.312$, $p = 0.046$<br>$t_9 = 5.051$, $p < 0.0007$<br>$t_9 = 9.398$, $p < 0.0001$<br>$t_9 = 13.38$, $p < 0.0001$<br>$t_9 = 20.00$, $p < 0.0001$<br>$t_9 = 9.435$, $p < 0.0001$<br>$t_9 = 4.098$, $p = 0.0027$<br>$t_9 = 9.685$, $p < 0.0001$<br>$t_9 = 15.18$, $p < 0.0001$<br>$t_9 = 13.40$, $p < 0.0007$<br>$t_9 = 15.18$, $p < 0.0001$<br>$t_9 = 6.646$, $p < 0.0001$<br>$t_9 = 7.561$, $p < 0.0001$<br>$t_9 = 5.669$, $p = 0.0003$<br>$t_9 = 6.578$, $p = 0.0001$<br>$t_9 = 3.275$, $p = 0.0096$<br>$t_9 = 8.822$, $p < 0.0001$ |
| 2a | Saline (n = 5)<br>D-amphetamine (n = 5) | Unpaired t-test (pS845-GluA1)<br>Vermis: $t_8 = 3.762$, $p = 0.0055$<br>Lateral: $t_6 = 3.811$, $p = 0.0089$<br>Unpaired t-test (pS831-GluA1)<br>Vermis: $t_8 = 0.4005$, $p = 0.69$<br>Lateral: $t_8 = 0.4376$, $p = 0.67$ |
| 2b | Saline (n = 4)<br>D-amphetamine (n = 4) | Unpaired t-test<br>pS845-GluA1: $t_6 = 2.649$, $p = 0.0381$<br>pS831-GluA1: $t_6 = 0.6776$, $p = 0.523$ |
| 2c | Saline (n = 7)<br>Methylphenidate (n = 7) | Unpaired t-test<br>pS845-GluA1: $t_{12} = 2.124$, $p = 0.055$<br>pS831-GluA1: $t_{12} = 2.624$, $p = 0.022$ |
| 3c | *Transcripts* (input, pellet)<br>*Slc6a3* : (n = 5, n = 5)<br>*Ddc* : (n = 5, n = 5)<br>*Th* : (n = 5, n = 5) | Unpaired t-test<br>$t_8 = 8.437$, $p < 0.0001$<br>$t_8 = 1.060$, $p = 0.32$<br>$t_8 = 1.394$, $p = 0.207$ |
| 4b | *Transcripts* (input, pellet)<br>**Drd1** : (n = 5, n = 5)<br>**Drd2** : (n = 5, n = 5)<br>**Drd3** : (n = 4, n = 5) | Unpaired t-test<br>$t_8 = 7.949$, $p < 0.0001$<br>$t_8 = 0.4887$, $p = 0.638$<br>$t_7 = 0.02035$, $p = 0.98$ |
| 4c | Saline-Saline (n = 5)<br>Saline-D-amphetamine (n = 6)<br>SCH23390-Saline (n = 6)<br>SCH23390- D-amphetamine (n = 6) | One-way ANOVA<br>$F_{(3, 19)} = 11.62$, $p = 0.0001$ |
| 4d | Saline (n = 6)<br>Cocaine (n = 6)<br>GBR12783 (n = 6) | One-way ANOVA<br>$F_{(2, 15)} = 0.9$, $p = 0.4143$ |
| 5c | *Transcripts* (control, VMAT2-cKO)<br>**Gria1** : (n = 7, n = 7)<br>**Drd1** : (n = 6, n = 7)<br>**Adrb1** : (n = 7, n = 7) | Unpaired t-test<br>$t_{12} = 0.8588$, $p = 0.407$<br>$t_{11} = 1.428$, $p = 0.181$<br>$t_{12} = 1.057$, $p = 0.311$ |
| 5d | Control Saline (n = 7)<br>Control D-amphetamine (n = 7)<br>VMAT2-cKO Saline (n = 7)<br>VMAT2-cKO D-amphetamine (n = 6) | Two-way ANOVA<br>Genotype $F_{(1, 23)} = 9.354$, $p = 0.0054$<br>Treatment $F_{(1, 23)} = 24.03$, $p < 0.0001$<br>Interaction $F_{(1, 23)} = 14.88$, $p = 0.0008$ |
| 5e | *Transcripts* (input, pellet)<br>**Adra1a** : (n = 6, n = 5)<br>**Adrb1** : (n = 6, n = 5)<br>**Adrb2** : (n = 5, n = 5) | Unpaired t-test<br>$t_9 = 8.176$, $p < 0.0001$<br>$t_9 = 9.276$, $p < 0.0001$<br>$t_8 = 0.6044$, $p = 0.567$ |
| 5g | Saline-Saline (n = 5)<br>Saline-D-amphetamine (n = 6)<br>Propranolol-Saline (n = 5)<br>Propranolol-D-amphetamine (n = 6)<br>Betaxolol-Saline (n = 5)<br>Betaxolol-D-amphetamine (n = 6) | One-way ANOVA<br>$F_{(5, 27)} = 8.904$, $p < 0.0001$ |
| 6a | Saline-Saline (n = 7)<br>Saline-D-amphetamine (n = 7)<br>Betaxolol-Saline (n = 6)<br>Betaxolol-D-amphetamine (n = 7) | One-way ANOVA<br>$F_{(3, 23)} = 8.128$, $p = 0.0007$ |
| 6b | *Transcripts* (saline, d-amphetamine)<br>**Adrb1** : (n = 6, n = 7)<br>**Adcy2** : (n = 10, n = 9)<br>**Ppp1r14c** : (n = 10, n = 9)<br>**Ppm1a** : (n = 10, n = 8)<br>**Pde7b** : (n = 10, n = 9) | Unpaired t-test<br>$t_{11} = 3.013$, $p = 0.0118$<br>$t_{17} = 2.347$, $p = 0.03$<br>$t_{17} = 5.203$, $p < 0.0001$<br>$t_{16} = 2.228$, $p = 0.04$<br>$t_{17} = 1.591$, $p = 0.13$ |
| 6c | Saline-Saline (n = 7)<br>Saline-Methylphenidate (n = 7)<br>Betaxolol-Saline (n = 7)<br>Betaxolol-Methylphenidate (n = 7) | One-way ANOVA<br>$F_{(3, 24)} = 23.71$, $p < 0.0001$ |
| 6d | *Transcripts* (saline, methylphenidate)<br>**Adrb1** : (n = 10, n = 10)<br>**Adcy2** : (n = 10, n = 9)<br>**Ppp1r14c** : (n = 10, n = 8) | Unpaired t-test<br>$t_{18} = 2.216$, $p = 0.047$<br>$t_{17} = 2.809$, $p = 0.0121$<br>$t_{16} = 3.552$, $p = 0.0027$ |
| | *Ppm1a* : (n = 10, n = 8)<br>*Pde7b* : (n = 10, n = 8) | $t_{16} = 2.858, p = 0.0114$<br>$t_{16} = 3.708, p = 0.0019$ |
| S4 | *Transcripts* (control, VMAT2-cKO)<br>*Adrb2* : (n = 7, n = 7)<br>*Adra1a* : (n = 7, n = 7)<br>*Ddc* : (n = 6, n = 7)<br>*Th* : (n = 6, n = 6)<br>*Drd3* : (n = 6, n = 6)<br>*Comt* : (n = 7, n = 7)<br>*Maoa* : (n = 7, n = 7)<br>*Maob* : (n = 7, n = 7)<br>*Drd2* : (n = 7, n = 7)<br>*Slc6a3* : (n = 6, n = 6)<br>*Gfap* : (n = 7, n = 7)<br>*Itgam* : (n = 7, n = 7)<br>*Aif1* : (n = 7, n = 7)<br>*Apq4* : (n = 7, n = 7)<br>*Kcnj10* : (n = 7, n = 7)<br>*Gria2* : (n = 7, n = 7) | Unpaired t-test<br>$t_{12} = 0.4212, p = 0.68$<br>$t_{12} = 0.7538, p = 0.465$<br>$t_{11} = 0.1486, p = 0.884$<br>$t_{10} = 1.306, p = 0.228$<br>$t_{10} = 0.8116, p = 0.436$<br>$t_{12} = 1.204, p = 0.25$<br>$t_{12} = 0.1638, p = 0.872$<br>$t_{12} = 1.151, p = 0.27$<br>$t_{12} = 0.1923, p = 0.85$<br>$t_{10} = 1.970, p = 0.077$<br>$t_{12} = 1.149, p = 0.162$<br>$t_{12} = 0.296, p = 0.7718$<br>$t_{12} = 0.416, p = 0.684$<br>$t_{12} = 0.208, p = 0.838$<br>$t_{12} = 0.6828, p = 0.507$<br>$t_{12} = 0.961, p = 0.355$ |
| S5 | *Transcripts* (input, pellet)<br>*Ppp1r14c*: (n = 9, n = 8)<br>*Ppm1a* : (n = 9, n = 8)<br>*Pde7b* : (n = 9, n = 8)<br>*Adcy2* : (n = 9, n = 8) | Unpaired t-test<br>$t_{15} = 8.638, p < 0.0001$<br>$t_{15} = 0.1095, p = 0.9143$<br>$t_{15} = 6.254, p < 0.0001$<br>$t_{15} = 13.02, p < 0.0001$ |

